# Differential Metabolite Production Underlies Disruption of the Cystic Fibrosis Airway Microbiota by Pathogens

**DOI:** 10.64898/2026.07.16.738945

**Authors:** Sydney M. Morabbi, Niladri Bhowmik, Shaz Sutherland, Evelyn A. Wylie, Reagan S. Decker, Akram Al Daerwish, Mercedes Pérez Pérez, Elizabeth Pascual, Erika I. Lutter, Benjamin Philmus, Reed M. Stubbendieck

**Author notes:** Address correspondence to Reed M. Stubbendieck.

## Abstract

Cystic fibrosis (CF) is a multisystem disease characterized by the accumulation of mucus in the airways that promotes pathogen colonization, leading to respiratory exacerbations, lung failure, and death. Culture-independent approaches have revealed that the CF airway harbors a complex microbiota, including opportunistic pathogens and bacteria that colonize the oropharynx. Here, we reanalyzed 5,260 16S rRNA gene microbiota datasets to infer ecological associations between members of the CF microbiota. We determined that pathogens are more likely to proliferate and dominate when present, while oropharyngeal bacteria are more likely to form persistent communities. Further, we found higher diversity and increasing numbers of inferred interactions were positively associated with lung function. In contrast, pathogens were negatively associated both with each other and with oropharyngeal bacteria, suggesting that they may disrupt the microbiota. To validate these predictions, we cultured 1,597 bacterial isolates from 96 people with CF and performed 12,542 coculture assays against eight representative CF pathogenic and oropharyngeal bacteria. 23% of these interactions resulted in growth inhibition. While *Pseudomonas* isolates were, on average, the most inhibitory, we observed variable activity among isolates. We then confirmed that *Pseudomonas aeruginosa* isolates, even those from the same donor and timepoint, exhibited significant differences in their metabolome and bioactivity profiles that correlated with acquisition of mutations. Together, our results suggest that pathogens may disrupt the CF microbiota and bloom in part through differential metabolite production. Furthermore, these data highlight that characterizing multiple isolates is necessary to capture the full landscape of chemically mediated interactions within microbial communities.

**Importance:** The cystic fibrosis (CF) airway harbors a complex microbiota, including oropharyngeal bacteria and opportunistic pathogens that establish chronic infections and cause lung failure. We confirmed that microbiota diversity is correlated with health and that a pathogen-dominated microbiota is associated with reduced lung function. We then inferred microbial interactions, which suggested that pathogens are able to disrupt the microbiota. To validate these predictions, we cultured bacterial isolates from people with CF and performed thousands of coculture assays, finding that approximately one-quarter of interactions resulted in growth inhibition. *Pseudomonas* broadly inhibited other members of the CF airway microbiota. However, we observed marked variability in bioactivity and metabolite profiles of *Pseudomonas aeruginosa* isolates, even from the same donor at the same time. Our results suggest that pathogens disrupt the CF microbiota, possibly through bioactive metabolite production, and that characterizing multiple isolates is necessary to capture the complete picture of interactions in these communities.

## Introduction

Cystic fibrosis (CF) is a genetic disease in humans that results from recessive mutations in the gene encoding the CF transmembrane conductance regulator. As a consequence of these mutations, chloride ion transport is impaired in the respiratory epithelium leading to the accumulation of a thick and dehydrated mucus layer in the lower airways (1, 2), which serves as the substrate for the microbial growth (3).

The most well-known bacterial pathogen that colonizes the airways of people with CF (pwCF) is *Pseudomonas aeruginosa.* A long-standing assumption in the field was that *P. aeruginosa* is the sole dominant pathogen in the CF airway. However, a recent meta-analysis revealed that less than half of sputum samples from 507 adults with CF (42.7%) are dominated by this pathogen (relative abundance >50%) (4). In addition to being colonized by other pathogens, such as *Burkholderia cenocepacia* and *Staphylococcus aureus* (5), recent advances in metagenomics and culturomics have confirmed that the airways in pwCF are colonized by a heterogeneous microbiota. This community can include other opportunistic pathogens (e.g., *Achromobacter, Burkholderia*, *Haemophilus, Pseudomonas,* and *Staphylococcus*) and commensal oropharyngeal bacteria (e.g., *Neisseria*, *Prevotella*, *Rothia*, *Streptococcus*, and *Veillonella*) (6, 7). Together, these ten highlighted genera were previously reported to represent >85% of reads identified by 16S rRNA gene amplicon sequencing of CF sputum (8, 9).

The microbiota of the CF airway is complex and the contribution of microbial interactions toward disease remains poorly understood (10, 11). Recent microbiome-wide association studies suggest that bacterial interactions are strong predictors of declining lung function in pwCF (12). For instance, coculture between CF pathogens can enhance virulence factor production or antibiotic resistance by one or both organisms (13). Further, anaerobic bacteria from the oropharynx can ferment mucins and supply *P. aeruginosa* with short-chain fatty acids as a carbon source (14). Disrupting these mucin fermenters predisposes *P. aeruginosa* to treatment with antibiotics that it normally resists (15, 16). In contrast, other studies report that oropharyngeal bacteria may be beneficial and that oral anaerobes are negatively correlated with CF pathogens. (7, 17). Therefore, competitive interactions in the CF airway may constrain the growth of specific microbes. However, it has also been suggested that elimination of specific pathogens in the CF airway may open niches for colonization by other opportunistic pathogens, which are normally suppressed by their competitors (9, 18). Thus, identifying the antagonistic interactions that structure the CF airway microbiota are therefore key to predicting disease trajectories and therapeutic outcomes.

Previous studies that investigated antagonism between CF bacteria have focused primarily on *P. aeruginosa* cocultured with *S. aureus* (19–21) and *Streptococcus* spp. (22). To date, there have been few large-scale interaction screens to identify antagonism between other members of the CF airway microbiota. For instance, Menetrey et al. recently performed 170 coculture interaction assays between CF and environmental isolates of *P. aeruginosa*, *S. aureus, Achromobacter xylosoxidans*, and *Stenotrophomonas maltophilia*, and documented 30 instances of growth inhibition, suggesting antagonistic interactions occur between among the CF airway microbiota (23). While their work was more exhaustive than prior studies, many members of the CF airway microbiota have not been tested in coculture with other microbes. Thus, there is a gap in our knowledge regarding antagonism that may occur between members of the CF airway microbiota, including interactions that could be exploited for human benefit.

One major mechanism by which bacteria mediate antagonism is through the production of bioactive secondary metabolites (24, 25). Our previous genomic surveys indicated that biosynthetic gene clusters for secondary metabolite are abundant in aerodigestive tract bacteria that reside within the airways of pwCF (26). Further, secondary metabolite production has been reported from CF pathogens, and these compounds function in a broad range of biological activities, including as antibiotics, siderophores, and virulence factors. For instance, *P. aeruginosa* produces a suite of secondary metabolites, including redox-active phenazines, such as pyocyanin, which contribute to host tissue damage and virulence (27–29), electron transport (30, 31), and microbial interactions (32–34). In addition, this pathogen also produces the iron-binding siderophores pyochelin and pyoverdine (35), and antibiotics, including the copper-containing fluopsin C (36), and 2-heptyl-4-hydroxyquinoline *N*-oxide (19, 37). However, given that *P. aeruginosa* populations diversify during chronic infection (38) and that secondary metabolite production often varies between strains (39–41), antagonistic phenotypes are unlikely to be uniform even within a single pwCF. This variability suggests that surveys that incorporate multiple isolates from the same pwCF and across multiple pwCF are needed to understand how bioactive metabolites mediate interactions broadly between members of the CF microbiota.

To address these gaps in the ecology of the CF airway, we combined a large-scale reanalysis of microbiota data with culture-based interaction screens. Using 5,260 existing 16S rRNA gene microbiota datasets, we found that pathogens and oropharyngeal bacteria exhibit differential colonization dynamics in the airways of pwCF. Further, we determined that lung function correlated positively with microbiota diversity and with the number of inferred interactions between bacterial genera. Among these potential associations, we ascertained that pathogenic genera exhibited mixed associations with other pathogens, but were negatively correlated with oral bacterial genera, all of which were positively correlated with each other. To test competitive interactions directly, we cultured 1,597 bacterial isolates from 96 pwCF and performed 12,542 coculture assays against eight CF-associated bacteria to assess their inhibitory activity. We found that CF isolates of *Pseudomonas*, *Bacillus, Klebsiella*, and *Stenotrophomonas* were the most inhibitory, while *Staphylococcus* and *Streptococcus* showed little inhibitory activity. However, we observed extensive variability in inhibitory activity of *Pseudomonas*, even among isolates from the same donor at the same timepoint. Subsequently, we generated extracts from a subset of these *P. aeruginosa* isolates and demonstrated that they exhibited variable metabolome and bioactivity profiles. We then identified mutations associated with BGCs or secondary metabolism broadly that were unique to a *P. aeruginosa* isolate that was less inhibitory in coculture. Together, these data suggest that microbial interactions contribute toward shaping the composition of the airway microbiota in pwCF and that isolate-level variability in bioactive metabolite production is a major and underappreciated factor underlying these interactions.

## Results

### Pathogens bloom while oropharyngeal bacteria form resident communities in the CF airway

We were initially interested in determining the effect of microbiota composition on lung function in pwCF. We reanalyzed 5,260 16S rRNA gene amplicon sequencing samples from 49 sequencing studies deposited in the Sequence Read Archive (SRA) (**Table S1**) using the same pipeline (see **Materials and Methods** for full details) and identified the composition of each microbiota. We limited our analysis to the genus level, as the amplicon sequencing data we analyzed used different regions of the 16S rRNA gene. Further, this marker possesses variable and low phylogenetic resolution below the genus-level (42, 43). This approach enabled us to consolidate all of the data into a single analysis, at the cost of losing lower level taxonomic resolution (i.e., amplicon sequence variants or operational taxonomic units). While this is a reasonable tradeoff, we are unable to differentiate between different *Achromobacter*, *Burkholderia*, *Staphylococcus*, and other genera where multiple species colonize the CF airway (44). Overall, there were 19 genera, including 7 genera with pathogens, present at ≥1% relative abundance (RA) in the meta-community of the airways of pwCF (**Table 1**).

**Table 1.** Composition of the bacterial meta-community of the CF airway. . All bacterial genera at ≥1.0% overall RA from 5,260 CF airway samples are shown. Pathogen-associated genera are indicated in **bold**, while oropharyngeal genera are indicated in plain text.

| Genus | Overall RA (%) | Mean RA (%) | Median RA (%) [IQR]<sup>a</sup> |
| --- | --- | --- | --- |
| *Streptococcus* | 23.2 | 22.2 | 15.1 [4.5-32.1] |
| ***Pseudomonas*** | **11.4** | **20.0** | **1.7 [0.1-31.3]** |
| ***Staphylococcus*** | **10.9** | **12.5** | **0.3 [0.0-11.2]** |
| *Prevotella* | 9.0 | 10.5 | 5.1 [0.6-15.7] |
| ***Haemophilus*** | **6.0** | **6.8** | **0.7 [0.0-4.6]** |
| *Veillonella* | 5.8 | 6.9 | 3.5 [0.6-10.2] |
| *Rothia* | 4.9 | 3.7 | 0.9 [0.1-4.4] |
| ***Stenotrophomonas*** | **2.9** | **6.0** | **0.0 [0.0-1.1]** |
| *Neisseria* | 2.9 | 3.3 | 0.3 [0.0-2.9] |
| *Granulicatella* | 2.4 | 2.6 | 0.9 [0.2-2.5] |
| ***Achromobacter*** | **2.3** | **6.1** | **0.0 [0.0-0.8]** |
| *Gemella* | 2.2 | 2.1 | 0.6 [0.1-2.6] |
| *Porphyromonas* | 1.9 | 3.1 | 0.4 [0.0-2.5] |
| ***Burkholderia*** | **1.8** | **6.6** | **0.0 [0.0-0.2]** |
| *Fusobacterium* | 1.2 | 2.4 | 0.2 [0.0-1.9] |
| *Actinomyces* | 1.1 | 0.8 | 0.2 [0.0-0.7] |
| *Schaalia* | 1.1 | 1.0 | 0.3 [0.1-1.0] |
| *Alloprevotella* | 1.0 | 1.4 | 0.2 [0.0-1.2] |
| ***Escherichia*** | **1.0** | **3.1** | **0.0 [0.0-0.2]** |
| Total | 93.0 |  |  |

<sup>a</sup>Interquartile range

The distribution pattern for each genus in the CF airway microbiota is highly skewed, indicating that subsets of the samples account for most of the abundance of each taxon. As one example, the mean RA of *Pseudomonas* is 20.0%, but its median RA is only 1.7% (interquartile range [IQR]: 0.1 to 31.3%) (**Table 1**), indicating that this genus tends to proliferate to high abundances when it is present.

Accordingly, *Pseudomonas* exhibits the highest dominance (defined as >50% RA within an individual sample) of any genus. In contrast, the rarer CF pathogenic genera (e.g., *Achromobacter*, *Burkholderia*, and *Stenotrophomonas*) tend to exhibit low prevalence (defined as ≥1% RA within an individual sample), but proportionally high bloom rates (defined as Dominance/Prevalence) (**Fig. 1A**). This pattern indicates that when these rarer CF pathogens are present, they tend to dominate the microbiota. The per-genus bloom rate of pathogens overall (median: 18.4%, IQR: 16.1 to 28.7%) is significantly higher compared to that of the oropharyngeal genera (median: 0.6%, IQR: 0.2 to 1.4%) (Wilcoxon rank-sum *W* = 1, *P* = 6.1e-4) (**Fig. 1A**). However, when different CF pathogens dominate the airways, they generally reach similar Ras (median dominant RA: 69.2 to 84.0%; Kruskal-Wallis χ^2^ = 17.6, df = 6, *P* = 0.007; all post-hoc Benjamini-Hochberg [BH]-adjusted *P* > 0.05, except for *Burkholderia* and *Stenotrophomonas* vs. *Escherichia*) (**Fig. S1A**). Furthermore, when pathogens dominate a sample, they reach higher RAs (median: 78.7%, IQR: 63.8 to 92.1%) compared to the oropharyngeal bacteria (median: 67.7%, IQR: 57.0 to 83.7%) (Wilcoxon rank-sum *W* = 528,157, *P* < 2.2e-16) (**Fig. S1B**). These patterns are consistent with pathogenic and oropharyngeal genera differing with respect to their colonization dynamics, suggesting that the latter form background resident communities, while the former proliferate when present.

**Figure 1.**
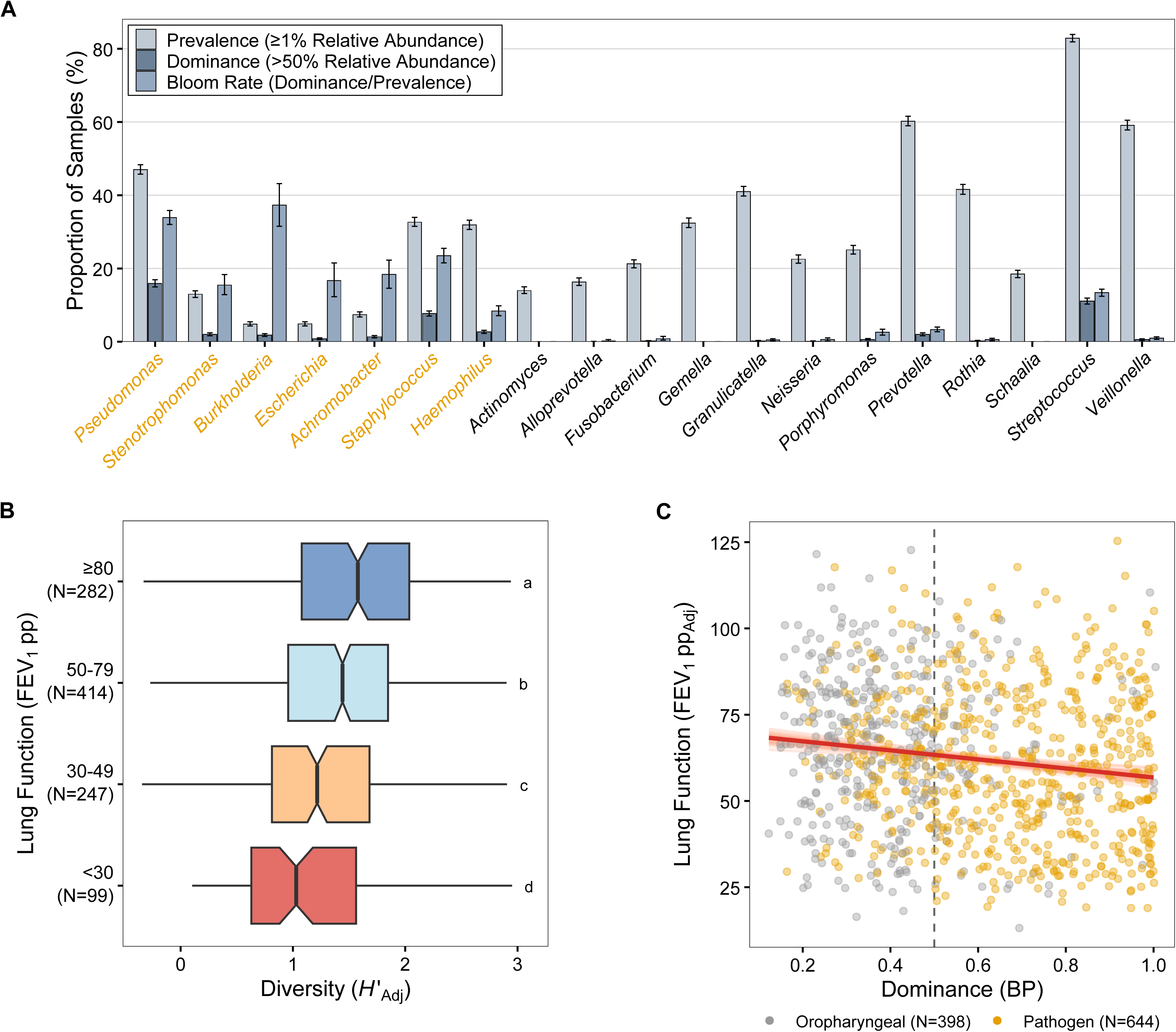
Loss of microbiota diversity and pathogen proliferation are associated with decreased CF lung function. (**A**) Bar plot showing the prevalence, dominance, and bloom rate for bacterial genera found in the airways of pwCF. The color of the bar indicates the reported statistic, as indicated by in the top-left. CF pathogenic strains are indicated in gold. (**B**) Box plots of the genus-level Shannon diversity index after adjustment for BioProject accession, age, and sample type (H′_Adj_) vs. lung function category based on the forced expiratory volume in 1 s percent predicted (FEV_1_ pp). Each microbiota sample is indicated with a black point. The upper and lower bounds of the box plots indicate the 75th and 25th percentiles, respectively. The horizontal black bars indicate the medians, and the notches represent the 95% confidence intervals of the medians. The whiskers extend from the bounds of the box to the largest and smallest values that are no further than ±1.5× the IQR. FEV_1_ categories that share letters are not significantly different (BH-adjusted *P* > 0.05). (**C**) Plot showing the relationship between the Berger-Parker (BP) Dominance and lung function after adjustment for BioProject accession, age, and sample type (FEV1 pp_adj_). Each microbiota sample is indicated with a point. Points were jittered to avoid overplotting. Gold points indicate samples where CF pathogens were the most abundant genera. Points to the right of the dashed line are dominated by a single genus. The red line indicates the line of best fit, and each of the 100 light red lines represents a bootstrap replicate of the fit. See **Figure S3** in the supplemental material for a version of this figure where the points are given different colors for each specific genus of CF pathogen.

### The composition of the CF airway microbiota is only weakly associated with age and lung function

Given the differences between the dynamics of pathogenic and oropharyngeal genera in the CF airway, we were then interested in determining if other factors influence the composition of this microbiota. We performed a permutational multivariate analysis of variance (PERMANOVA) analysis and determined that BioProject accession (F_48,5260_ = 33.61, R^2^ = 0.24, BH-adjusted *P* = 1.2e-4), 16S rRNA gene region sequenced (F_7,5260_ = 35.17, R^2^ = 0.045, BH-adjusted *P* = 1.2e-4), and sample type (F_5,5260_ = 33.1, R^2^ = 0.031, BH-adjusted *P* = 1.2e-4) were the primary contributors to variation in microbiota composition (F_53,5260_ = 17.66, R^2^ = 0.15, BH-adjusted *P* = 1.2e-4) at the genus-level, indicating that observed differences were largely driven by technical factors. Subsequently, we performed partial PERMANOVA analyses on subsets of the samples conditioned on these factors and found statistically significant, but weak effects of age category (F_4,2386_ = 2.13, R^2^ = 0.0031, BH-adjusted *P* = 7.4e-4) (**Fig. S2A**) and lung function category, as measured by the forced expiratory volume in 1 s percent predicted (FEV_1_ pp) (F_3,1103_ = 2.67, R^2^ = 0.0063, BH-adjusted *P* = 2.3e-4) (**Fig. S2B**), but not sex (F_1,1618_ = 0.96, R^2^ = 0.0005, BH-adjusted *P* = 0.45) (**Fig. S2C**), on the microbiota composition. However, the age and FEV_1_ categories only reached statistical significance in 19.1% and 12.3%, respectively, of 1,000 random subsets of 100 samples. This subset analysis indicates that their statistical significance is driven by the large sample size and minimally by biological effects. Taken together, these analyses suggest that age, sex, and lung function exert little-to-no effect on the composition of the CF airway microbiota at the genus-level.

### Decreasing microbiota diversity and increasing pathogen dominance are associated with decreased lung function in CF

We next examined if other microbial factors were related to lung function. We determined the alpha diversity of CF airway samples, as estimated with the Shannon diversity index (H′), was positively associated with lung function in pwCF (Kruskal-Wallis χ^2^ = 42.41, df = 3, *P* = 3.3e-9; values after adjustment for age, BioProject accession, 16S rRNA region sequenced, and sample type) (**Fig. 1B**), consistent with prior reports (6, 45). We then wanted to characterize the effect of microbiota dominance on lung function. We determined dominance for each sample using the Berger-Parker index (BP), which represents the proportion of the microbiota occupied by the most abundant genus. We observed an inverse correlation between dominance and lung function (regression coefficient [b] = −1.4 FEV_1_ pp per 0.1-unit increase in BP, *t* = −4.9, *P* = 8.9e-7; full-model adjusted R^2^ = 0.21, with covariates of age, BioProject accession, 16S rRNA region sequenced, and sample type) (**Fig. 1C**), consistent with a prior report (45). Notably, the majority (86%) of CF airway samples with a BP value ≥0.5, indicating numeric dominance by a single genus, are pathogen-dominated (**Fig. 1C, Fig. S3**). Together, these data are consistent with the hypothesis that loss of microbiota diversity and proliferation of pathogens contributes to worsening disease in the airways of pwCF (45).

### Negative associations involving pathogens and positive associations among oropharyngeal bacteria correspond to divergent lung function outcomes in CF

We then wanted to explore potential causes of pathogen blooms in the CF airway and their consequence on lung function. It was recently reported that increasing interactions between CF pathogens are associated with decreasing lung function (12). In contrast to this result, we observed that as the number of predicted interacting genera in a sample increased, lung function improved (b = +1.4 FEV_1_ pp per genus, *t* = 6.9, *P* = 9.5e-12; full-model adjusted R^2^ = 0.24, with covariates of age, BioProject accession, 16S rRNA region sequenced, and sample type) (**Fig. 2A**). To place this finding in the context of the microbiota diversity analyses from above, we compared the relative contributions of interactions, dominance, and diversity to lung function.

**Figure 2.**
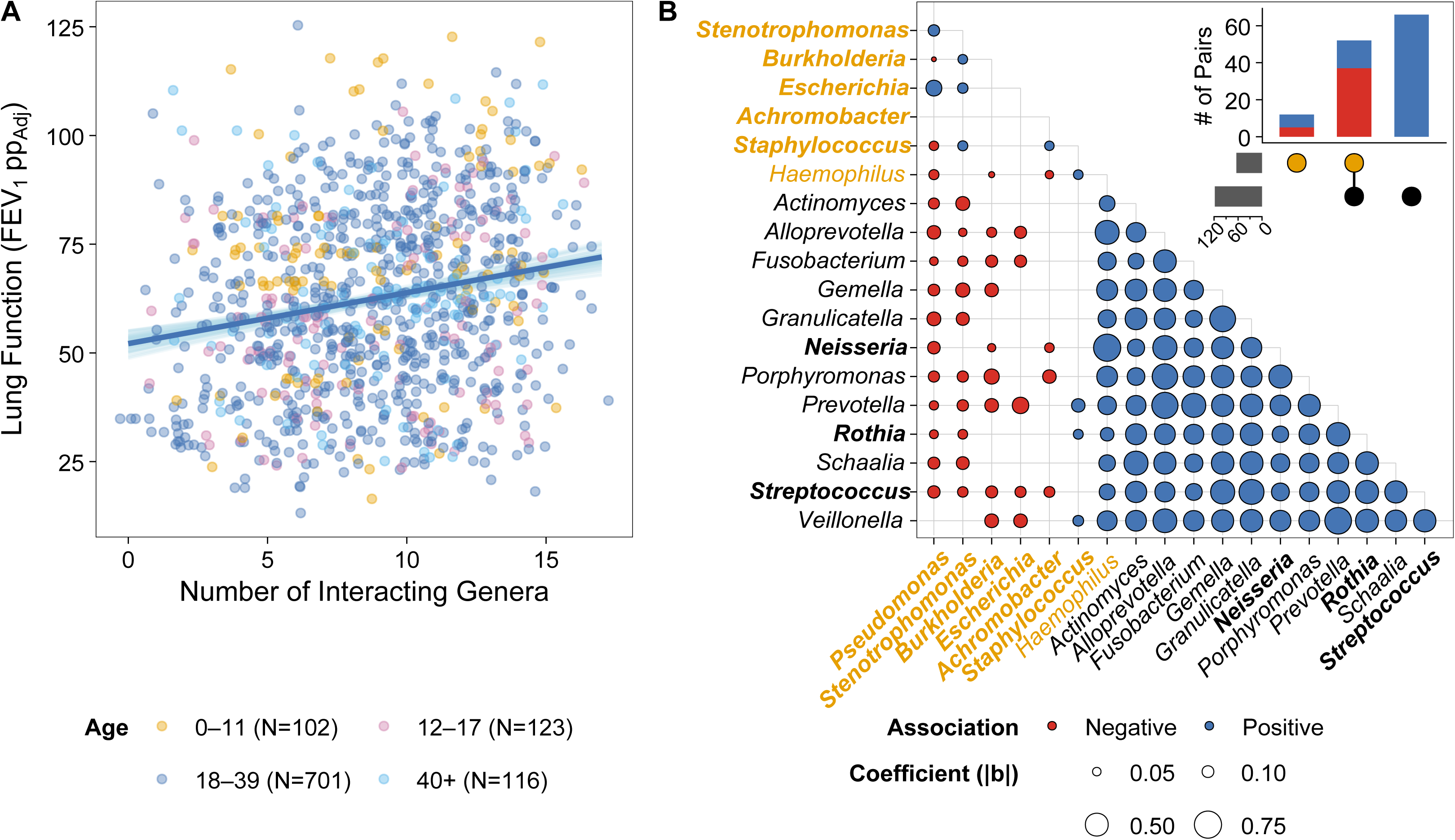
**Increasing bacterial interactions are associated with increased CF lung function**. (**A**) Plot showing the relationship between the number of inferred interacting genera and lung function after adjustment for age, BioProject accession, 16S rRNA gene region sequenced, and sample type (FEV_1_ pp_Adj_). Each microbiota sample is indicated with a dot, and the color corresponds to the age category indicated below. Points were jittered to avoid overplotting. The blue line indicates the line of best fit, and each of the 100 light blue lines represents a bootstrap replicate of the fit. (**B**) Bubble plot showing associations between bacterial genera in the CF airway based on 5,260 16S rRNA gene amplicon sequencing studies. Genera with CF pathogens are indicated in gold. For genera indicated in bold, there was at least one corresponding CF isolate or target included in coculture assays. Red and blue bubbles indicate negative and positive associations, respectively, and the size of the bubble indicates the strength of the association as measured by the absolute value of the regression coefficient (|b|). For simplicity, only the lower half of the correlation matrix is shown. Inset: UpSet plot showing the number of significant association pairs by category (top-right). The gold and black dots represent pathogens and oropharyngeal bacteria, respectively. (bottom-left) Bar plot indicating the total number of associations that involve pathogens and oropharyngeal bacteria on the top and bottom, respectively.

Dominance had the strongest effect (absolute value of the standardized regression coefficient [|β*|] = 3.45 FEV_1_ pp/standard deviation [SD]), followed by the number of inferred interactions (|β*| = 3.21 FEV_1_ pp/SD), and microbiota diversity (|β*| = 2.35 FEV_1_ pp/SD). However, it is important to note here that each of these factors are interrelated.

We then used linear models to evaluate positive and negative correlations between the abundances of different bacterial genera and infer potential microbial interactions that occur within the CF airway. Of 12 significant associations that occurred between two pathogen-containing genera, 7 (58.3%) were positive and 5 (41.7%) were negative. Notably, in 52 significant associations between pathogenic genera and oropharyngeal bacteria, 15 (28.8%) were positive and 37 (71.2%) were negative. Most of the positive associations occurred between *Haemophilus* or *Staphylococcus* and oropharyngeal bacteria (**Fig. 2B**). In contrast, we found that all 66 significant associations between oropharyngeal bacteria were positive (**Fig. 2B**). In summary, these microbiota data indicate that interactions within the CF airway are linked to lung health. Specifically, these data suggest that while oropharyngeal bacteria exhibit potential cooperative interactions and are thought to be beneficial for CF lung function (17), pathogens exhibit antagonistic interactions and disrupt the microbiota.

### Isolation and identification of bacteria from CF clinical samples

We wanted to test if the negative associations that we inferred from amplicon sequencing studies would replicate in the laboratory. In prior work, sputa samples were obtained from pediatric and adult donors with CF at the Oklahoma City Cystic Fibrosis Clinic and cultured aerobically on non-selective medium (e.g., brain-heart-infusion [BHI] or skim milk [SM] agar plates). Each plate was scaped and frozen as pools of bacterial colonies. We then generated a collection of CF bacterial isolates by culturing these pools on a panel of ten differential and selective media (see **Materials and Methods** for full detail on sample collection and media composition). In total, we obtained 1,597 isolates from 96 pwCF. Specific donor and bacterial isolate details are provided in **Table S2** and **Table S3**, respectively, in the supplementary material.

To identify the CF isolates, we performed colony PCR and high-throughput sequencing of the 16S rRNA gene V3-V4 region and then classified sequences using the eHOMD (46). Together, the genera *Staphylococcus* (n=630), *Pseudomonas* (n=558), *Bacillus* (n=88), *Enterococcus* (n=78), *Lacticaseibacillus* (n=53), *Klebsiella* (n=38), *Streptococcus* (n=38), *Escherichia* (n=19), *Stenotrophomonas* (n=19), and *Microbacterium* (n=12) represented 96% of the total isolates that we cultured (**Table S3**). Note, each of these genera were detected in the amplicon sequencing data, though some were below 1% mean RA. Given their abundance in pwCF, high growth rate, and non-fastidious nature, it is not surprising that *Pseudomonas* and *Staphylococcus* were the most common isolates in our collection. The remaining 4% of isolates (n=64) consist of genera with <10 representative isolates or were unclassified at the genus level. We classified these latter isolates as “Other” (**Table S3**). Together, this collection of isolates contains bacterial genera representing 7 of 19 of the most abundant genera (≥1% RA) in the CF meta-community and 7 of 14 of the most abundant (≥1% RA) aerobic or facultative anerobic genera (**Table 1**).

### CF isolates inhibit respiratory bacteria

We next wanted to test our predictions for antagonistic interactions among the CF microbiota by assessing the ability of these 1,597 CF isolates to inhibit a single strain of the following bacteria: *A. xylosoxidans*, *B. cenocepacia*, *Burkholderia cepacia*, *Neisseria meningitidis*, *P. aeruginosa*, *Rothia mucilaginosa*, *S. aureus*, and *Streptococcus oralis* using the cross-streak method. We chose these bacteria as targets because they represented a combination of known pathogens and commensal organisms that commonly colonize the CF airway (7, 9), and were amenable to culturing in our laboratory. We chose *N. meningitidis* as a representative *Neisseria,* and as a pathogen of concern, and *S. oralis* as a representative *Streptococcus* that grew adequately under our culture conditions. Exclusion of a representative *Prevotella* spp., *Veillonella* spp., or other obligate anaerobic bacterial strains was intentional due to the technical challenge of performing coculture assays with a combination of strictly aerobic and anaerobic target organisms.

We cultured all 1,597 CF isolates on BHI agar plates. After allowing the cultures to establish overnight, we inoculated the targets perpendicularly to the CF isolate. After overnight incubation, we measured the zone of inhibition (ZOI), which we defined as the distance between the isolate and first growth of the target streaks (**Fig. 3A**). In total, we measured ZOIs from 12,542 total isolate × target combinations. In the case of 234 of these combinations, there was poor growth of the target on a monoculture control plate, overgrowth of the CF isolate, or contamination. We did not consider these combinations in further analysis. As 44 out of 64 isolates in the “Other” category did not inhibit any of the targets tested, we did not consider any isolates in this category further in this study. We assigned each isolate × target combination a score based on the ZOI of the target: no inhibition (ZOI: 0 cm, score: 0), weak inhibition (ZOI: 0 to 1.0 cm, score: 1), moderate inhibition (ZOI: 1.0 to 2.0 cm, score: 2), strong inhibition (ZOI: ≥2.0 cm or no observed growth of target, score: 3).

**Figure 3.**
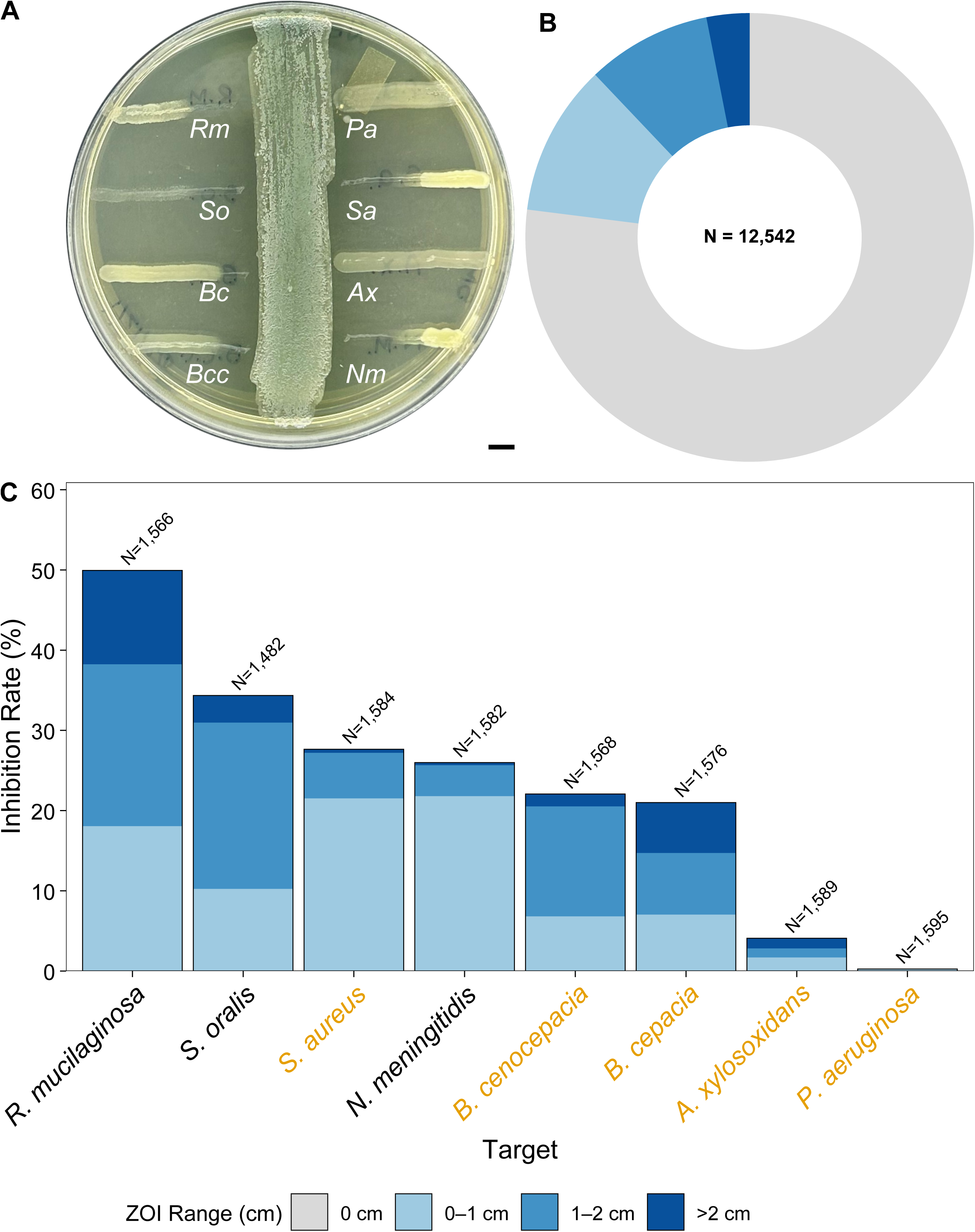
CF isolates inhibit the growth of respiratory target strains. (**A**) Photograph of a representative coculture between *Pseudomonas* isolate NB0851 (middle of plate) and eight bacterial target strains. The ZOI is defined as the distance between the CF isolate and the point of growth on the perpendicular target streak. Scale bar = 5 mm. Abbreviations: *Ax*, *A. xylosoxidans* subsp. *xylosoxidans*; *Bc*, *B. cepacia; Bcc*, *B. cenocepacia*; *Nm*, *N. meningitidis*; *Pa*, *P. aeruginosa*; *Rm*, *R. mucilaginosa*; *Sa*, *S. aureus*; *So*, *S. oralis* subsp. *oralis*. Scale bar = 5 mm. (**B**) Donut plot showing the fraction of isolate × target interactions with each color representing no inhibition (ZOI: 0 cm, score: 0), weak inhibition (ZOI: 0 to 1.0 cm), moderate inhibition (ZOI: 1.0 to 2.0 cm), strong inhibition (ZOI: ≥2.0 cm or no observed growth of target), as indicated by the legend below. The total number of tested interactions is indicated in the center of the donut. (**C**) Stacked bar plot showing the frequency of inhibition for each target strain. Each color represents the proportion of interactions that resulted in the corresponding level of inhibition, as indicated by the legend below. The total number of interactions tested involving a given target is specified above each stacked bar. CF pathogens are indicated in gold.

In total, we observed that 2,886 (23% of the total) isolate × target combinations resulted in some level of growth inhibition of the target (i.e., ZOI: >0 cm, score: 1 to 3) (**Fig. 3B**). There were significant differences in inhibition of each target (χ^2^ = 2489.9; df = 21; Monte Carlo simulated *P* < 1.0e-5). We found that *R. mucilaginosa* was the most susceptible to inhibition (50% of interactions resulted in inhibition), followed by *S. oralis* (34%). We observed similar inhibition of *B. cepacia, B. cenocepacia*, *N. meningitidis*, and *S. aureus* (21 to 28%). As expected, *A. xylosoxidans* (4%) and *P. aeruginosa* (0.3%) were the least inhibited targets (**Fig. 3C**). In the latter case, there were only four isolates that inhibited *P. aeruginosa*, albeit very weakly (average ZOI: 0.1 cm). We identified one of these inhibitory isolates as a *Microbacterium* and the other three isolates as *Pseudomonas,* which were all derived from the same donor and may represent the same strain.

We then sought to determine which CF genera were most inhibitory toward the selected targets by normalizing the inhibition exhibited by each genus by the total number of coculture assay combinations tested. There were significant differences in target inhibition based on isolate genus (χ^2^ = 2612.3; df = 27; Monte Carlo simulated *P* < 1.0e-5). Consistent with the inferred associations (**Fig. 2C**), *Pseudomonas* was the most inhibitory genus that we tested (47% of combinations resulted in inhibition) and was approximately 1.5× more inhibitory than the next most inhibitory genus. *Stenotrophomonas*, *Bacillus*, and *Klebsiella* were modestly inhibitory genera (∼30%). *Enterococcus* and *Escherichia* were poorly inhibitory genera (∼16%). We observed little inhibition by *Microbacterium*, *Lacticaseibacillus*, *Staphylococcus*, and *Streptococcus* (<10%) (**Fig. 4A**). In the latter case, lack of inhibition by *Staphylococcus* and *Streptococcus* was expected based on the inferred associations (**Fig. 2C**).

**Figure 4.**
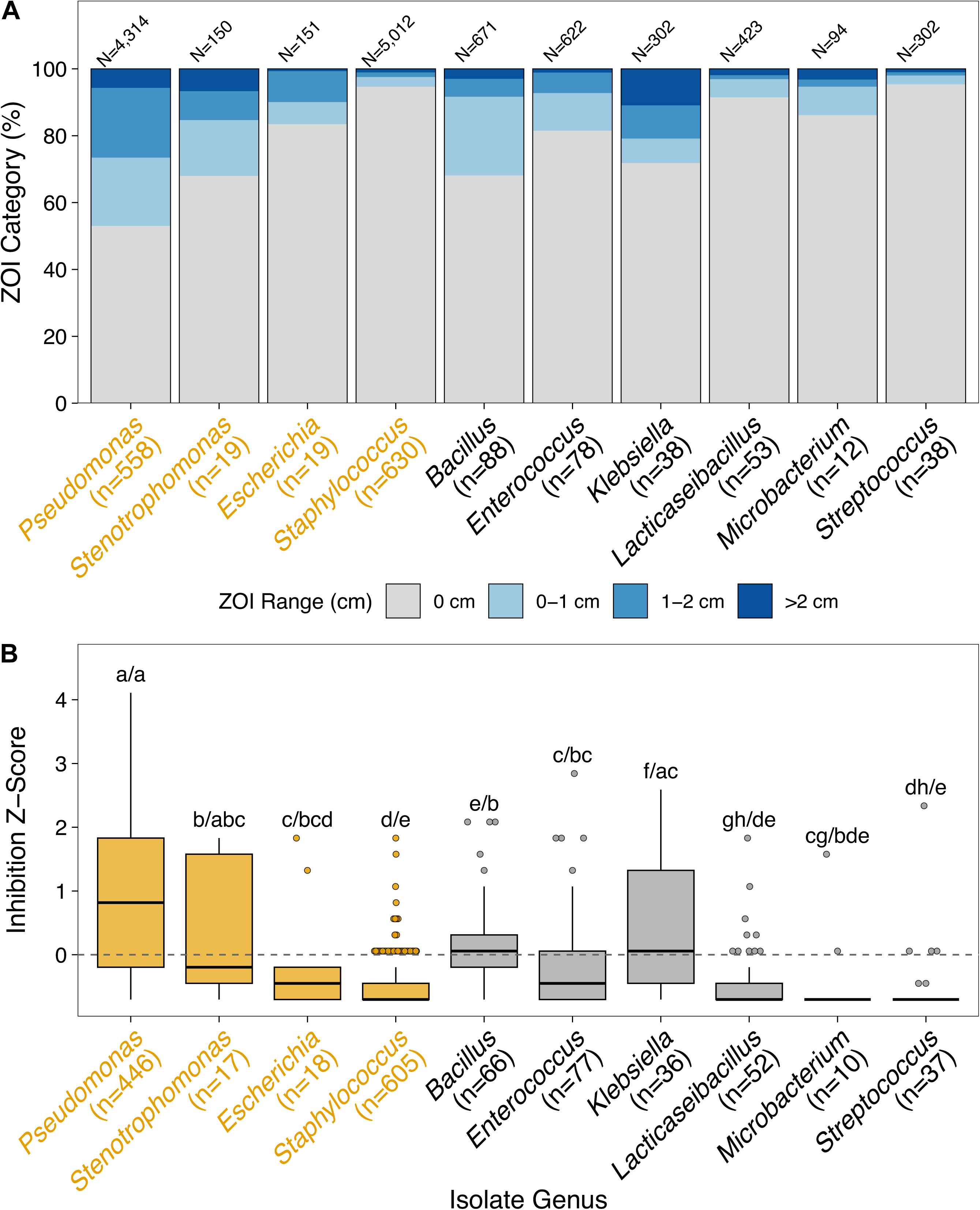
CF isolates exhibit variable inhibition of respiratory target strains. **(A)** Stacked bar plot showing the frequency of inhibition exhibited by each genus of CF isolates tested. Each color represents the proportion of interactions that resulted in the corresponding level of inhibition of a respiratory target strain, as indicated by the legend below. The total number of isolates tested (n), and the total number of interactions assayed (N), involving a given CF isolate genus are specified below and above each stacked bar, respectively. (**B**) Plot showing the Z-score normalized inhibition of each CF isolate tested against all respiratory target strains. The upper and lower bounds of the box plots indicate the 75th and 25^th^ percentiles, respectively. The horizontal black bars indicate the medians. The whiskers extend from the bounds of the box to the largest and smallest values that are no further than ±1.5× the IQR. Outliers are shown as points. The dashed line indicates the mean inhibition of all isolates (i.e., Z-score = 0). The first letter corresponds to the comparisons in Z-scores between genera and the second letter after the slash (/) corresponds to the Z-score dispersion between genera. Genera that share letters are not significantly different (BH-adjusted *P* > 0.05). CF pathogens are indicated in gold.

### CF isolates possess variable bioactivity against respiratory bacteria

We wanted to identify individual isolates that exhibited strong inhibitory activity. To compare the overall inhibitory activity spectrum of each individual CF isolate, we calculated and compared Z-scores based on the total inhibition score across the eight targets. As before, we observed significant differences when comparing the median inhibition Z-scores between different genera (Kruskal-Wallis χ^2^ = 514.65; df = 9; *P* < 2.2e-16). (**Fig. 4B**). Isolates of *Pseudomonas* tended to be the most inhibitory, while isolates of *Staphylococcus* andn *Streptococcus* were the least inhibitory. Isolates of *Bacillus*, *Klebsiella*, *Enterococcus*, *Escherichia*, *Microbacterium*, and *Lacticaseibacillus* exhibited intermediate inhibition (**Fig. 4B**). In addition, across the focal genera, there were significant differences in Z-score dispersion, indicating intra-genus variability with respect to inhibitory activity (Fligner-Killeen χ^2^ = 527.36; df = 9; *P* < 2.2e-16). Subsequently, we identified 23 significant differences between the Z-score distribution of genera pairs, with *Pseudomonas* exhibiting the most significant differences when compared to the other genera (**Fig. 4B**).

### Metabolome profiles vary between *Pseudomonas* isolates from the same donor that exhibit different inhibitory activity

Our coculture inhibition data indicated that *Pseudomonas* is the most antagonistic bacterial genus found in the CF airway. However, there was also significant variability among inhibitory activity of *Pseudomonas* isolates, with some isolates exhibiting strong inhibition and others that were less inhibitory. (**Fig. 4B**). Further, we identified 17 cases in which *Pseudomonas* isolates obtained from the same donor at the same timepoint displayed variable inhibitory activity (within-genus Z-score spread: ≥2) (**Fig. 5** and **Fig. S4** in the supplemental material). These observations suggested the potential of within-host diversification of isolates leading to changes in antagonistic interactions between *Pseudomonas* and other members of the CF microbiota.

**Figure 5.**
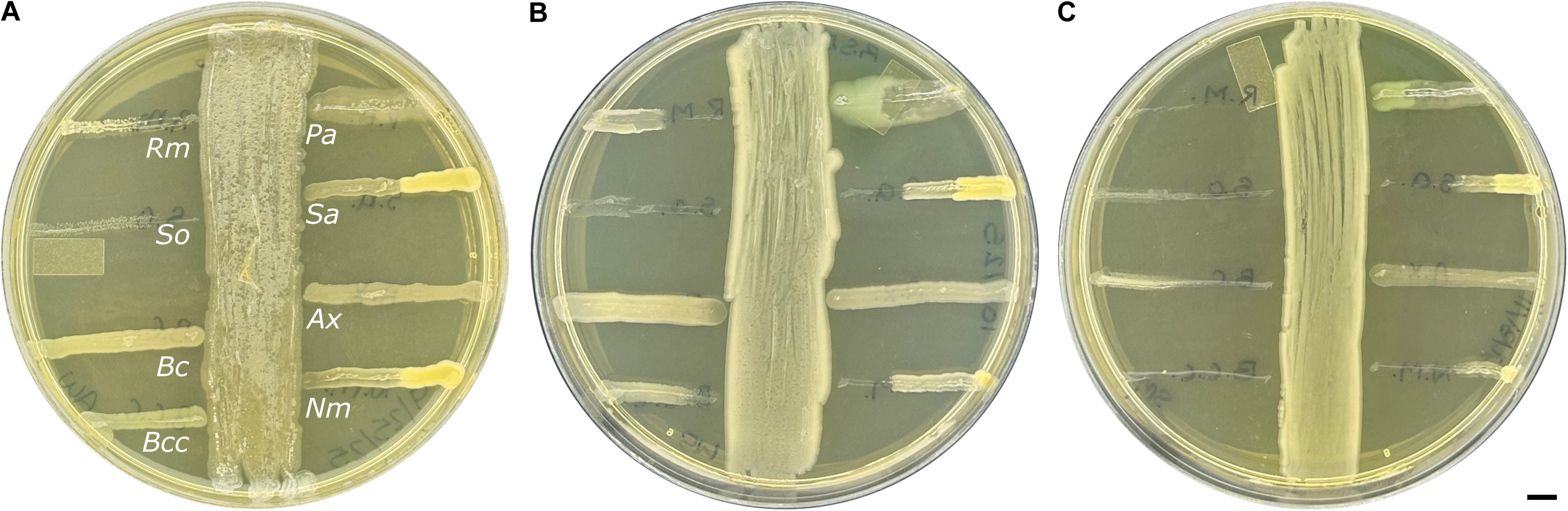
*Pseudomonas* isolates from the same donor and timepoint exhibit variable inhibition of respiratory target strains. Photograph of a coculture between *P. aeruginosa* isolate (**A**) NB1386 (Z-Score: −0.86), (**B**) RSD0018 (Z-Score: 0), and (**C**) NB0219 (Z-Score: 1.72) and eight target strains. Scale bar = 5 mm. Abbreviations: *Ax*, *A. xylosoxidans* subsp. *xylosoxidans*; *Bc*, *B. cepacia; Bcc*, *B. cenocepacia*; *Nm*, *N. meningitidis*; *Pa*, *P. aeruginosa*; *Rm*, *R. mucilaginosa*; *Sa*, *S. aureus*; *So*, *S. oralis* subsp. *oralis*.

We hypothesized that differences in bioactive metabolite produced by these *Pseudomonas* isolates were responsible for their variable inhibitory activity and could contribute toward their ability to displace other members of the microbiota. To test this hypothesis, we selected two donors (HID07 and HID08) from whom we obtained *Pseudomonas* isolates with varying inhibitory activity in cross-streak assays. We cultured isolates with low, medium, and high Z-scores in triplicate and then assessed the bioactivity of their cell-free supernatants (CFS) against *S. aureus* as a representative respiratory target.

The CFS of inhibitory *Pseudomonas* isolates (Z-score ≥ 0) inhibited *S. aureus* growth, while CFS from the less-inhibitory isolate NB1386 (Z-Score: −0.86) had no effect on *S. aureus* growth (see **Fig. S5A** in the supplemental material). In contrast, CFS from isolate NB1046 (Z-score: −0.43) did inhibit *S. aureus*. However, this isolate was also active against *S. aureus* in the coculture assay.

To determine if there were differences in the composition of CFS from inhibitory and less-inhibitory *Pseudomonas* isolates, we performed untargeted liquid chromatography coupled to tandem mass spectrometry (LC-MS/MS). In total, we aligned 5,645 molecular features (i.e., an *m*/*z* at a specific retention time), of which 468 varied significantly (multiclass Kruskal-Wallis *P* ≤ 0.01) between the six isolates. These molecular features spanned the full chromatographic run and mass range (**Fig. 6A**). We then wanted to assess whether the metabolome profiles clustered based on the donor or the inhibitory activity. We hierarchically clustered the 468 significant features, which split the CFS from *Pseudomonas* into two well-supported groups (**Fig. 6B**). We performed marginal PERMANOVA, which indicated that the donor significantly explained variation between *Pseudomonas* isolates (ω^2^ = 0.31, *P* = 0.011), whereas inhibitory phenotype had no effect (ω^2^ = 0.20, *P* = 0.078) (**Fig. S5B**). Together, these data suggest within-host diversification of *P. aeruginosa* metabolism occurs and that variation in metabolome profiles could be responsible for differing inhibitory activity of these pathogens.

**Figure 6.**
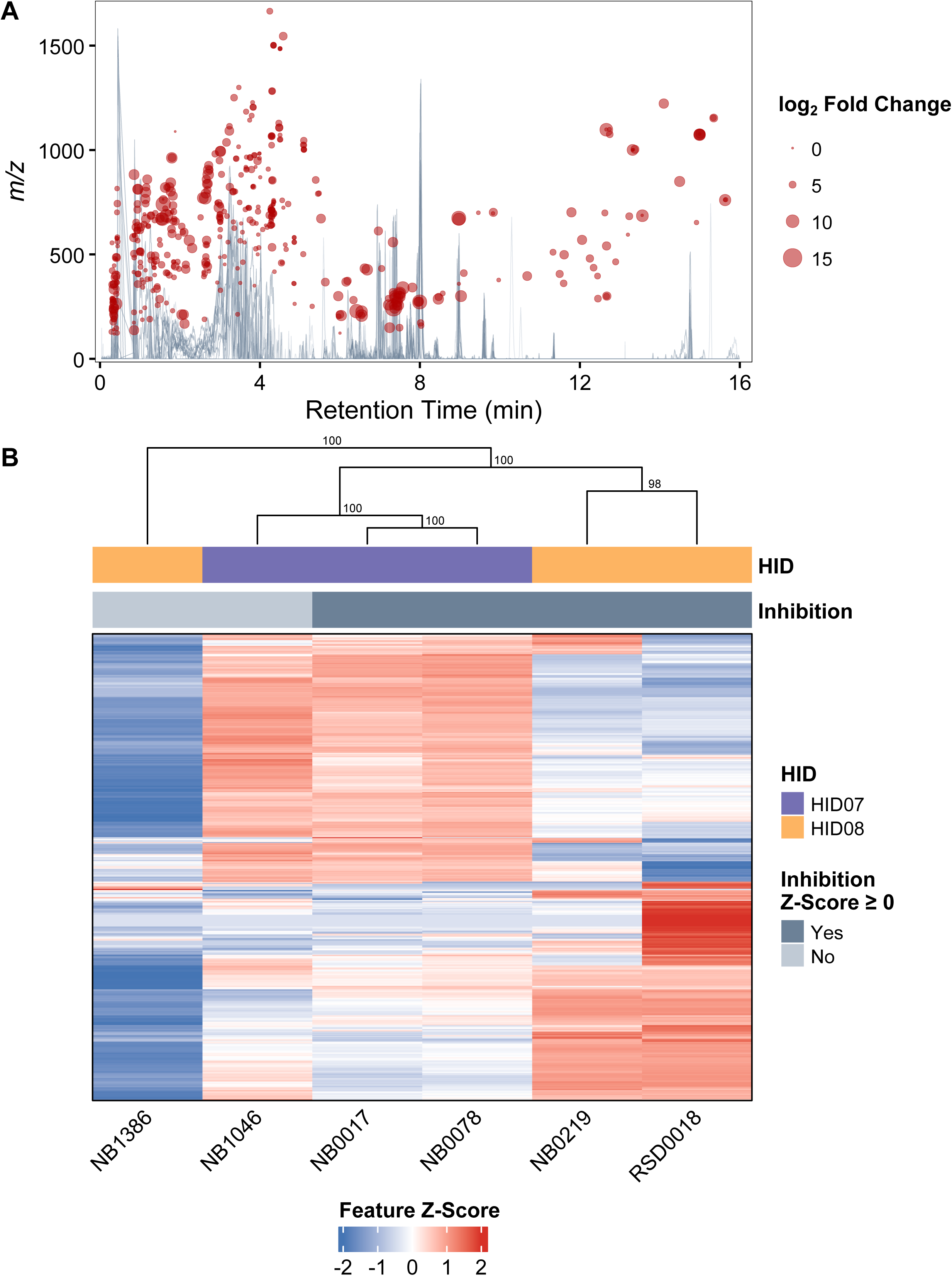
**The metabolomes of inhibitory and less-inhibitory *Pseudomonas* isolates are distinct**. (**A**) Cloud plot comparing the metabolome profiles of clinical *Pseudomonas* isolates. Traces represent the total ion chromatograms (TICs) of each of six clinical *Pseudomonas* isolates in triplicate after background subtraction of the mean medium blank. The dark blue and gray TICs are from extracts of inhibitory and less-inhibitory *Pseudomonas* isolates, respectively. Bubbles indicate 468 LC-MS/MS features that differ significantly (Kruskal-Wallis *P* ≤ 0.01) across different clinical *Pseudomonas* isolates. The size and shade of each bubble correspond to the log_2_ fold change and −log_10_ *P*-value, respectively, as indicated on the right. (**B**) Heatmap of the 468 significant features (rows) across the 6 isolates (columns). The feature intensities were averaged, log_2_-transformed, per-feature Z-scored, as indicated by the scale below. The isolates were clustered using Euclidean distance and the numbers at each node of the dendrogram represent bootstrap support (pvclust, 1,000 bootstraps). The top color bars represent the donor identity (HID) and inhibitory phenotype, as indicated on the right.

### Genetic differences between inhibitory and less-inhibitory *P. aeruginosa* isolates explain differences in metabolome profiles

We next wanted to identify specific genetic differences between *Pseudomonas* isolates that could explain the variation that we observed in their inhibitory activity and metabolome profiles. As mentioned previously, the metabolomes of the three isolates from HID08 grouped based on inhibition, as expected. In contrast, the metabolome of NB1046 from HID07 grouped with those of the inhibitory isolates, despite being classified as a less-inhibitory isolate, based on the coculture assay. We generated closed genomes using long-read sequencing for each of these six isolates and confirmed that they were all *P. aeruginosa* (average nucleotide identity [ANI] to the type-strain *P. aeruginosa* ATCC 10145^T^: 99.3 to 99.4%). We further determined that each set of three isolates from the same donor likely represent clonal lineages (ANI ≥ 99.995%), but were distinct between donors (ANI: 99.32%).

We then predicted the secondary metabolite biosynthetic gene cluster (BGC) content of each *P. aeruginosa* isolate genome using antiSMASH. Despite the isolates exhibiting variable inhibitory activity in coculture, they all encoded the same 17 BGCs, though two of the BGCs (16 and 17) were found in variable chromosomal positions between isolates from the different donors (**Fig. 7**). This finding prompted us to ask if the differences between these isolates could be explained by single-nucleotide polymorphisms (SNPs), insertions or deletions (indels), or other mutations in the inactive isolates. We identified differences between the isolates and mapped them back to the genome. In total, we only identified 7 variants (2 SNPs and 5 indels) between the three isolates from donor HID07 (**Fig. 7A**, **Table S4** in the supplemental material), consistent with more similarity between their metabolomes (**Fig. 6B**). Specifically, we identified three variants that were unique to isolate NB1046, consisting of a Pro41fs mutation in *mpl* encoding the UDP-MurNAc:L-Ala-γ-D-Glu-meso-DAP ligase, a W275R mutation in a gene encoding a DUF2875-containing protein, and a SNP in an intergenic region. We do not currently understand how these variants could affect the inhibitory activity of this isolate. In contrast, we identified 73 variants (55 SNPs, 17 indels, and 1 mixed SNP/Indel) between the three isolates from donor HID08, of which 62 were unique to the less-inhibitory NB1386 (**Fig. 7B**). Notably, isolate NB1386 contained multiple mutations in the BGCs for the siderophores pyoverdine and pyochelin (see **Table S5** in the supplemental material).

**Figure 7.**
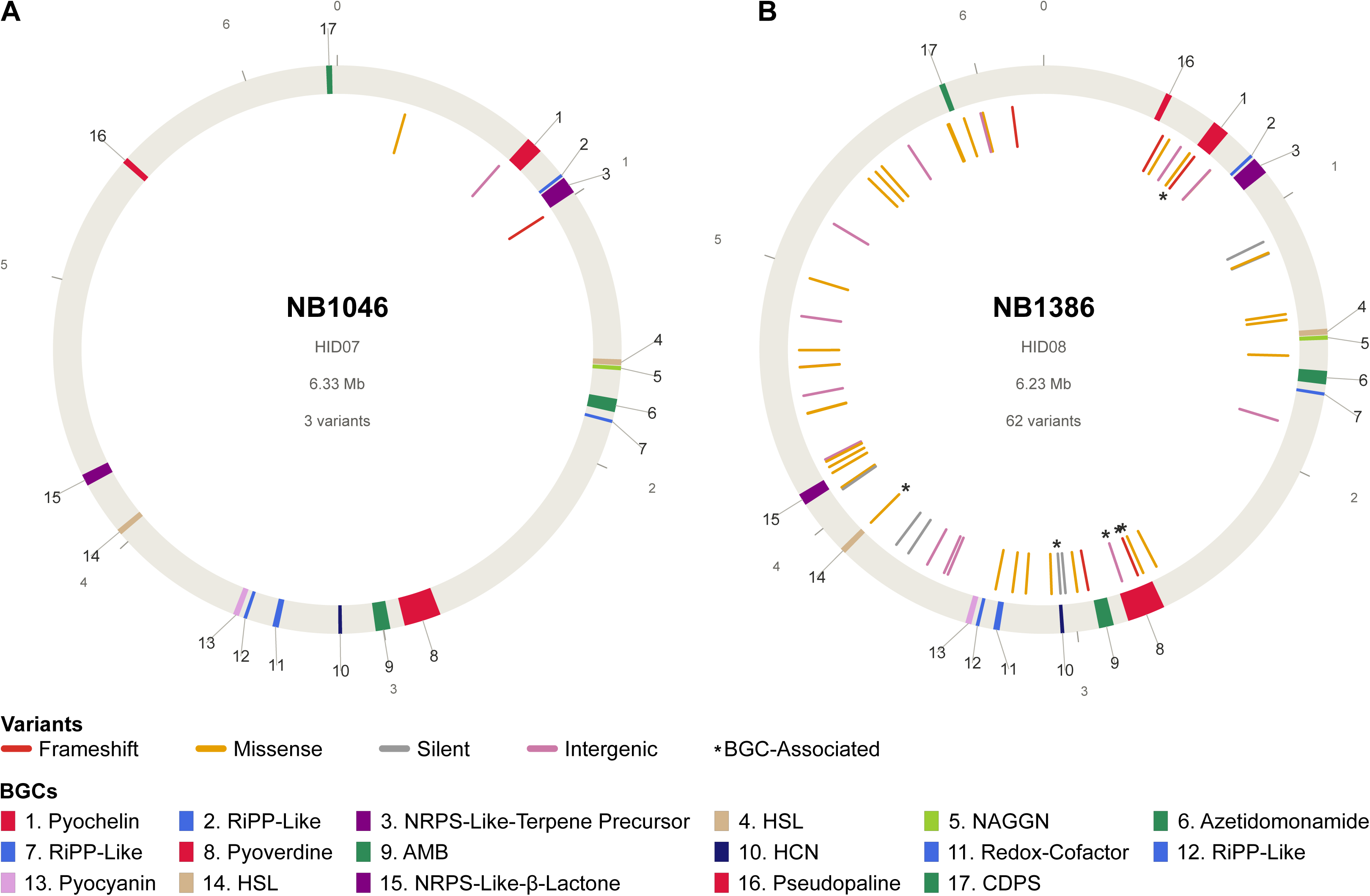
Genomic variation in less-inhibitory *P. aeruginosa* isolates tracks with changes metabolome profiles. Genome maps of the less-inhibitory *P. aeruginosa* isolates (**A**) NB1046 and (**B**) NB1386. The outer gray ring represents the genome with *dnaA* at the 0 position. The light gray numbers on the exterior of the circle indicate chromosomal position in Mb. Ticks on the interior of the circle represent genetic variants in the genome relative to inhibitory isolates from the same donor and their color represents the type of variant, as indicated by the legend below. BGCs are indicated with color strips and numbered, as indicated by the legend below. Variants that occur within a BGC are marked with an asterisk. Abbreviations: RIPP, ribosomally synthesized and post-translationally modified peptide; NRPS, non-ribosomal peptide synthetase; HSL, homoserine lactone; NAGGN, N-acetylglutaminylglutamine amide; AMB, L-2-amino-4-methoxy-trans-3-butenoic acid; HCN, hydrogen cyanide; CDPS, tRNA-dependent cyclodipeptide synthase.

In addition, the less-inhibitory isolate NB1386 has multiple additional mutations outside of BGCs that could affect secondary metabolism. For instance, the start codon of *lasI*, which encodes the synthase for the quorum sensing molecule *N*-3-oxododecanoyl homoserine lactone (3-oxo-C12-HSL) is mutated to an Ile codon, likely abrogating protein translation. Furthermore, *relA* harbors an in-frame deletion and a missense mutation, and *spoT* has a frame shift, together suggesting loss of the stringent response (see **Table S5** in the supplemental material). Both quorum sensing and the stringent response have been previously demonstrated to regulate secondary metabolite production in *P. aeruginosa* (47–49). Together, these data suggest that loss of inhibitory activity in isolate NB1386 and other *Pseudomonas* isolates may be due, in part, to mutations in genes that are essential for secondary metabolite biosynthesis or growth rate control.

## Discussion

In this study, we investigated the impact of microbial interactions on microbiota dynamics and disease in the airways of pwCF. We initially analyzed preexisting 16S rRNA gene amplicon sequencing datasets, which led us to determine that members of the oropharyngeal microbiota and pathogens exhibit different colonization dynamics within the airways of pwCF (**Fig. 1**). We then inferred interactions that occur between these two different groups of bacteria and found that increasing numbers of predicted interactions were associated with lung function (**Fig. 2**). To validate the negative associations that we inferred, we performed culture from CF airway specimens to build an isolate library, which we then used for *in vitro* coculture inhibition assays with ecologically relevant bacterial strains. We confirmed that antagonism was widespread among CF isolates (**Fig. 3** and **4**). We then noticed that different isolates of *Pseudomonas* possessed variable bioactivity in coculture (**Fig. 4** and **5**). Finally, we showed that different *P. aeruginosa* isolates, even those from the same person at the same time point, produced different metabolites and exhibited variable bioactivity *in vitro* (**Fig. 6** and **7)**.

Previous studies have reported a well-defined ecological succession of bacterial populations that occurs within the airways of pwCF as they age (50). In infancy the airways are first colonized primarily by bacteria from the oropharynx and upper respiratory tract (e.g., *Streptococcus* spp. and *Haemophilus influenzae*). Then in childhood and adolescence the abundance of *S. aureus* rises. Finally, *P. aeruginosa* or other, rarer pathogens proliferate in adulthood and as disease advances (6). In our reanalysis of 16S rRNA gene amplicon sequencing studies, we detected age-associated shifts in the composition of the microbiota, albeit only weakly (**Fig. S2**). However, our data were consistent with a broader ecological pattern described in prior work (6, 51), as oropharyngeal bacteria tended to form relatively resident communities, whereas pathogens tended to bloom in abundance and dominate the microbiota when they are present (6, 45, 51). These pathogen blooms were associated with loss of lung function, supporting the idea that pathogen proliferation is linked to disease progression. Alternatively, this pattern is also consistent with the possibility that the inflammatory and nutrient-rich environment of the CF airway selects for pathogens that are adapted to grow under these conditions (52, 53). Regardless, the association between pathogen blooms and loss of lung function suggests that disruption of the microbiota, whether by environmental changes, host immunity, or direct microbial interactions, contributes to disease progression in pwCF.

Reanalysis of 16S rRNA gene amplicon sequencing studies suggested the potential for widespread antagonistic and cooperative interactions among members of the CF airway microbiota (**Fig. 2**). Notably, we found that increasing numbers of interacting genera were associated with improved lung function in pwCF (**Fig. 2A**). This observation appears to contrast with Rivett et al. (12), who recently reported that increasing numbers of interactions among a subset of CF-associated taxa were associated with worsening disease in the CF airway. However, in our analysis, many of the inferred interactions among oropharyngeal genera were positive, and these bacteria are typically associated with a more diverse microbiota and with lung health (7, 17). Rivett et al. instead focused on taxa whose pairwise associations were linked to reduced lung function, including pathogens. These studies suggest that positive associations among oropharyngeal bacteria are characteristic of resident communities in healthier pwCF, whereas worsening disease is associated with antagonistic interactions involving pathogens that disrupt the microbiota. This interpretation is strengthened by our coculture assays, which showed that *Pseudomonas* and other pathogens inhibit the growth of other members of the CF airway microbiota (**Fig. 4**), consistent with prior reports that tested smaller numbers of isolates (23). Taken together, these results support a model in which pathogen blooms in the CF airway result, at least in part, from antagonistic interactions that disrupt the oropharyngeal microbiota and destabilize the community.

Comparison of the metabolomes (**Fig. 6**) and genomes (**Fig. 7**) of inhibitory and less-inhibitory *P. aeruginosa* isolates suggests that production of bioactive metabolites is at least, in part, responsible for the ability of these pathogens to antagonize other members of the CF airway microbiota. Notably, several *P. aeruginosa* secondary metabolites have been reported to reach concentrations between 1 and 100 *µ*M in sputum from pwCF (54–56). These concentrations are sufficient to influence the growth of other microbes. We suspect that other CF pathogens may also produce secondary metabolites that contribute to disruption of the microbiota, but beyond several examples from *Burkholderia* spp. (57–66), not much is known about the compounds produced by other pathogens. It is important to note that the mechanisms responsible for disruption of the airway microbiota by *P. aeruginosa* and other CF pathogens are likely multifaceted and probably also involve production of secreted enzymes, type VI secretion systems, and other effectors (24). Further, antagonism may not be the only means used by pathogens to become dominant. For example, *Staphylococcus* blooms in approximately 25% of samples it is detected in (**Fig. 1A**), yet isolates of *Staphylococcus* were not predicted to be negatively associated with other bacteria (**Fig. 2B**), and these isolates were largely non-inhibitory (**Fig. 4**). Whether dominance of the microbiota by *Staphylococcus* is due solely to priority effects stemming from its early colonization or are due to alternative mechanisms will require additional investigation.

The finding of *P. aeruginosa* isolates with variable inhibitory activity from the same donor at the same timepoint (**Fig. 5-7**) highlights the broad diversification that can occur within pwCF over the course of chronic infection. These observations are consistent with several prior studies that have identified diversification of *P. aeruginosa* that occurs longitudinally within the CF airway (67, 68). In fact, production of the secondary metabolites pyocyanin and pyoverdine has been reported to be lost in *P. aeruginosa* over the course of chronic infection, possible as an adaptation to surviving in the environment of the CF airway (69, 70). Our results also indicate that these mutations can have consequences for microbial interactions that occur within this environment. Notably, most culture-independent approaches would fail to capture the differences that we observed. This observation demonstrates the importance of characterizing multiple isolates from either host or environmental samples to determine how they interact with the host or other members of the community and should be applied more broadly to culture-dependent surveys of other microbial ecosystems.

Here, we note there were limitations in our study. First, due to prior sample preparation protocols that did not limit exposure to oxygen, we were unable to culture strict anaerobes from the donated sputa samples. We predicted multiple interactions that involve strictly anaerobic bacteria (**Fig. 2B**) and it is known that anaerobes play important roles in interactions with CF pathogens (14, 15). In addition, we did not isolate any *Burkholderia* spp., as no sputa was donated from individuals at the Oklahoma City Cystic Fibrosis Clinic that were culture-positive for these pathogens. Second, we have focused exclusively on the bacteria that colonize the airways of pwCF, while ignoring *Aspergillus*, *Scedosporium*, and other fungi that often inhabit this same environment. These fungi are known interact with CF bacteria (18) and to produce bioactive metabolites (71–76). It is also likely that fungi will exhibit isolate-to-isolate variability in their bioactive metabolite production. Finally, our cross-streak assays are not well-suited to test for positive interactions (e.g., growth promotion) between microbes. While these limitations mean that our cultures are not representative of the overall CF microbiota, to our knowledge they do represent the largest study of interactions between CF microbes to date, and we recapitulate the associations we inferred from amplicon sequencing data. Future studies that account for each of these limitations will improve our understanding of the microbial interactions that occur within the CF airway.

In conclusion, we report that *P. aeruginosa* and other CF pathogens potentially disrupt the structure of microbiota, bloom in abundance, and cause worsening disease in part by producing bioactive metabolites, an underappreciated phenomenon in the CF airway. We speculate that molecules that target secondary metabolite production by these pathogens could be developed as an anti-virulence strategy to keep pathogen proliferation in check and prevent worsening lung disease in pwCF.

Furthermore, we suspect that additional efforts to screen the CF airway microbiota for bioactive metabolites may yield additional leads that could be developed as new antibiotics.

## Materials and Methods

### Microbiota analyses

We identified 16S rRNA gene amplicon sequencing studies from pwCF by searching the SRA for “cystic fibrosis” AND “16S”, followed by manual curation of the resulting studies and cross-validation with the corresponding publications to limit our scope to relevant specimens (i.e., airway, bronchoalveolar lavage, bronchoscope, oropharynx, protected brush, sputum) (12, 77–113). In total, we analyzed 5,260 sequencing runs (see **Table S1** in the supplemental material for the full list of runs analyzed and the donor demographics). For all analyses, we treated each individual sequencing run as an independent sample.

We downloaded reads from the SRA using fasterq-dump 2.10.3 from the SRA toolkit (https://github.com/ncbi/sra-tools). We then used fastp 0.23.2 (114) for quality control and adapter trimming of the raw reads. To remove potential contamination from human-derived reads, we used bowtie2 2.4.1 and discarded reads that mapped to a reference human genome sequence (GenBank accession: GCA_009914755.4). We then used mothur 1.48.1 (115) to process 16S rRNA gene amplicon sequencing reads and perform taxonomical classification using the human oral microbiome database (HOMD) 16S rRNA RefSeq V15.23 as the reference (https://www.homd.org//ftp/16S_rRNA_refseq/HOMD_16S_rRNA_RefSeq/V15.23/) (46). We determined the beta diversity by computing Bray-Curtis dissimilarities using the parallelDist package, followed by non-metric multidimensional scaling (NMDS) using the vegan package (116) in R. To test whether microbiota composition differed by host or technical variables, we performed PERMANOVA with the vegan package using 9,999 permutations. We tested these variables with both unadjusted and partial models conditioned on the technical covariates (BioProject accession, 16S rRNA gene region sequenced, and sample type). We adjusted all PERMANOVA *P*-values for multiple comparisons using the BH false discovery rate. To assess the effect of sample size on statistical significance, we performed 1,000 subsamples of 100 samples and reran PERMANOVA with 999 permutations per replicate.

We calculated the H′ for each sample using the vegan package. To remove any confounding effects of age, BioProject accession, 16S rRNA gene region sequenced, or sample type, we adjusted H′ using an ordinary least-squares model. We defined H′_Adj_ as the residuals from this model plus the overall mean H′. We excluded any sample from this model that was missing one or more of these covariates. We determined differences in the alpha diversity across FEV_1_ pp severity categories using the Kruskal-Wallis rank-sum test and performed post-hoc pairwise comparisons using Wilcoxon rank-sum tests using the BH false discovery rate. We calculated BP for each sample as the proportional abundance of the single most abundant genus. We considered samples to be pathogen-dominated if the combined RA of the genera *Achromobacter*, *Burkholderia*, *Escherichia*, *Haemophilus*, *Pseudomonas*, *Staphylococcus*, and *Stenotrophomonas* was ≥50% RA. We removed confounding effects of technical factors on BP, as above, for the statistical model.

### Interaction analysis

To characterize potential interactions between genera in the CF airway microbiota, we applied pairwise linear regression as previously described (12). For these analyses, we only included genera with meta-community RA ≥0.5% and a prevalence of ≥5% across all samples. Briefly, we determined all possible pairwise linear regressions between the abundances of two genera (denoted g_1_ and g_2_) by fitting the ordinary least squares model: log_10_(g_1_+1) ∼ log_10_(g_2_+1) and using the regression slope (b_x_) as a measure of the interaction directionality (negative or positive). We assessed significance using a Bonferroni-corrected threshold of α = 0.05/number of genus pairs tested. To identify pairs of genera whose co-abundance interacted to predict lung function, we fit a linear model for each genus pair using the following model: FEV_1_ pp ∼ log_10_(g_1_+1) + log_10_(g_2_+1) + (log_10_(g_1_+1) × log_10_(g_2_+1)). We determined the interaction term coefficient (b_i_) and *P*-value for each interaction pair and corrected for multiple hypothesis testing using the BH FDR. We designated all genera appearing in at least one significant pair as interaction-associated. For each sample, we calculated n_g_ as the count of interaction-associated genera detected (reads >0). We then tested the association between n_g_ and lung function while controlling for confounders by fitting a multiple regression: FEV_1_ pp ∼ n_g_ + Age + BioProject Accession + 16S rRNA gene region sequenced + Sample Type. We adjusted each FEV_1_ pp for these confounders using the residuals plus the mean, as above. We generated bootstrap confidence bands from 100 resampled trend lines.

### Isolation and maintenance of bacteria from CF samples

Sputum and oropharyngeal samples from 96 pediatric and adult pwCF at the Oklahoma Cystic Fibrosis Clinic were previously collected and stored in the laboratory of Dr. Erika Lutter (see **Table S2** in the supplementary material for donor information). This study was deemed “not human subjects research” by the University of Oklahoma Institutional Review Board (IRB). Sputum samples were collected from patients by the clinical staff in 50 mL sterile conical tubes. The clinic staff provided information on the patient age, gender, and medical condition upon admittance to the clinic, and a deidentified number was assigned to each sample. These samples were then cultured on non-selective BHI (0.6% [wt/vol] brain heart infusion, 0.6% [wt/vol] peptic digest of animal tissue, 0.5% sodium chloride [NaCl], 0.3% [wt/vol] dextrose, 1.45% [wt/vol] pancreatic digest of gelatin, 0.25% [wt/vol] disodium phosphate, 1.5% [wt/vol] agar) or SM (10% [wt/vol] skim milk, 1.5% [wt/vol] agar) plates aerobically at 37°C for 3 days. Afterwards, these plates were scraped, and all bacterial biomass was cryopreserved at −80°C in 10% (wt/vol) skim milk. We generated an isolate library by streaking these samples onto the following differential and selective media at 37°C for approximately one week: *Burkholderia cepacia* Selective Agar (BCSA; 1.0% [wt/vol] casein peptone, 1.0% [wt/vol] lactose, 1.0% [wt/vol] sucrose, 0.5% [wt/vol] sodium chloride, 226 *µ*M phenol red, 1.5% [wt/vol] agar, with the following additions after autoclaving: 5 *µ*M crystal violet, 10 *µ*g/mL gentamicin, 2.5 *µ*g/mL vancomycin, 60 *µ*g/mL polymyxin B, pH 7.0); Cetrimide Agar (CA; 2.0% [wt/vol] gelysate peptone, 0.14% [wt/vol] magnesium chloride, 1.0% [wt/vol] potassium sulfate, 0.03% [wt/vol] cetrimide, 1% [vol/vol] glycerol, 1.5% [wt/vol] agar, pH 7.2); Eosin Methylene Blue (EMB; 1.0% [wt/vol] gelatin peptone, 1.0% [wt/vol] lactose, 0.2% [wt/vol] dipotassium hydrogen phosphate, 0.04% [wt/vol] eosin yellow, 0.0065% [wt/vol] methylene blue, 1.5% [wt/vol] agar, pH 7.1); *Haemophilus* Isolation Agar (HIA; 1.6% [wt/vol] casein peptone, 0.5% [wt/vol] sodium chloride, 0.5% [wt/vol] protease peptone, 0.8% [wt/vol] BHI, 0.2% [wt/vol] dextrose, 0.25% [wt/vol] disodium phosphate, 1.5% [wt/vol] agar, with the following additions after autoclaving: 109 *µ*M nicotinamide adenine dinucleotide, ≥8,250 U bacitracin, 10 mg hemin, and 5.0% [vol/vol] defibrinated horse blood, pH 7.3); LBVT.SNR (1.0% [wt/vol] tryptone, 0.5% [wt/vol] yeast extract, 0.5% [wt/vol] sodium chloride, 1 mM sodium hydroxide, 86 *µ*M neutral red, 1.5% [wt/vol] agar, with the following additions after autoclaving: 1.0% wt/vol sucrose, 3 *µ*g/mL vancomycin, and 3 *µ*g/mL trimethoprim); Mannitol Soy Agar (MSA; 1.0% [wt/vol] protease peptone, 0.1% [wt/vol] beef extract, 7.5% [wt/vol] sodium chloride, 1.0% [wt/vol] D-mannitol, 70 *µ*M phenol red, and 1.5% [wt/vol] agar, pH 7.4), MacConkey media (1.7% [wt/vol] gelatin peptone, 1.0% [wt/vol], sodium chloride, 0.15% [wt/vol] bile salts, 0.15% [wt/vol] casein peptone, 0.15% [wt/vol] meat peptone, 104 *µ*M neutral red, 2.5 *µ*M crystal violet, 1.35% [wt/vol] agar, pH 7.1) supplemented with 0.5% [wt/vol] lactose (MCL) or 0.5% [wt/vol] xylose with the following additions after autoclaving: 20 *µ*g/mL vancomycin, 20 *µ*g/mL aztreonam, and 5 *µ*g/mL amphotericin B (MCXVAA); Mitis-Salivarius agar with Tellurite (MSAT; 5.0% [wt/vol] sucrose, 1.0% [wt/vol] proteose peptone, 1.0% [wt/vol] tryptose, 0.4% [wt/vol] dipotassium phosphate, 0.1% [wt/vol] dextrose, 78 *µ*M trypan blue, 2.0 *µ*M crystal violet, 1.5% [wt/vol] agar, with the following addition after autoclaving: 0.001% [wt/vol] potassium tellurite, pH 7.0); *Rothia* Enrichment Medium agar (REM; 2.5% [wt/vol] heart infusion, 1.5% [wt/vol] agar, with the following additions after autoclaving: 10 *µ*g/mL colistin and 0.5 *µ*g/mL lincomycin). We used GasPak jars containing carbon dioxide generators (BD), as necessary, for anaerobic incubation. We selected ≥2 colonies of each distinct morphology per sample per media type and passaged the strains aerobically on BHI agar plates to obtain pure cultures. We preserved all isolates at − 80°C in 25% (vol/vol) glycerol. We provide details for all bacterial isolates used in this study in **Table S3** in the Supplemental Material. For general propagation and manipulation, we cultured bacterial isolates on BHI agar plates or in BHI broth aerobically at 37°C with shaking.

### Identification of bacterial isolates

Identification of all bacterial isolates was performed via colony PCR and high-throughput sequencing in 96-sample batches at the Oklahoma State University Microbiomics and Culturomics Core Facility. Briefly, single bacterial colonies on BHI agar plates were touched with a pipette tip and resuspended in 20 *µ*L sterile, nuclease-free water at heated at 95°C for 10 min. The 16S rRNA gene V3-V4 region was amplified using primers 341F (5′-[Illumina Adapter]-[6 nucleotide randomized tag]-CCTACGGGNGGCWGCAG-3′) and 805R (5′-[Illumina Adapter]-[6 nucleotide randomized tag]-GACTACHVGGGTATCTAATCC-3′). Each 25-*µ*L reaction mixture contained 3 *µ*L lysate, 20 pmol of each primer, and 12.5 *µ*L of DreamTaq 2× Master Mix (ThermoScientific) with the following PCR cycling conditions: 95°C for 2 min; 25 cycles of 95°C for 30 s, 55°C for 30 s, and 72°C for 30 s; and a final extension at 72°C for 5 min. PCR amplification was confirmed for all samples using 1.0% (wt/vol) agarose tris-acetate-EDTA (TAE) gels.

Subsequently, a Research Silicon-A plate kit (Zymo Research) was used for PCR cleanup following manufacturer’s instructions. Nextera XT DNA library preparation kits (Illumina) were used for library preparation. Each 25-*µ*L reaction mixture contained 5 *µ*L of cleaned PCR product, 2.5 *µ*L i7, 2.5 *µ*L i5, and 12.5 *µ*L 2× master mix with the following PCR cycling conditions: 95°C for 3 min; 8 cycles of 95°C for 30 s, 55°C for 30 s, and 72°C for 30 s; and a final extension at 72°C for 5 min. As before, PCR amplification was confirmed using TAE gels. These samples were then cleaned using a ratio of 1:0.6 volume sample to AMPure XP magnetic beads (Beckman Coulter). The beads and samples were incubated for 5 to 8 min at ambient temperature, washed twice with 200 *µ*L of 80% (vol/vol) ethanol, air dried for 3 to 5 min, and eluted with 50 *µ*L of 10 mM Tris-HCl, pH 8.5. The libraries were quantified using a Qubit dsDNA HS fluorometer (Invitrogen) and normalized to equimolar concentration and pooled per 384-sample batch. The libraries were then sequenced using a NextSeq 2000 P1 600 Cycle (2×300 bp), following the platform loading guidance.

The pooled sequencing reads were demultiplexed into per-colony FASTQ files and basic read processing was performed using mothur, as above. We used the classify.seqs function in mothur to identify isolates using the eHOMD 16S rRNA RefSq V15.23 (46) with an 80% bootstrap confidence cutoff. If we failed to obtain a PCR product from the high-throughput colony PCR method, we then performed targeted colony PCR and Sanger sequencing. The identities of these isolates are provided in **Table S3** in the Supplemental Material.

### Cross-streak coculture assays

We inoculated 3 mL BHI broth with single colonies of CF bacterial isolates and incubated the cultures at 37°C overnight with shaking. We then spread 20 *µ*L of the undiluted overnight culture in a ∼8 × ∼1.5 cm-wide line in the center of a 25-mL BHI agar plate (diameter: 8.6 cm). We incubated the plates aerobically for 18 to 20 h at 37°C. We prepared overnight (16 to 18 h) cultures of *A. xylosoxidans* subsp. *xylosoxidans* strain DSM 2402, *B. cenocepacia* strain K56-2, *B. cepacia* strain DSM 7288, *N. meningitidis* strain DSM 10036, *P. aeruginosa* strain PID25, *R. mucilaginosa* strain HM-1055, *S. aureus* strain PID40 by inoculating single colonies into 3 mL BHI broth and incubating cultures at 37°C with shaking. We prepared cultures of *S. oralis* subsp. *oralis* strain DSM 20627 in tryptone soy broth (TSB; 1.7% [wt/vol] pancreatic digest of casein, 0.3% [wt/vol] papaic digest of soybean, 0.25% [wt/vol] dextrose, 0.5% [wt/vol] NaCl, 0.25% [wt/vol] dipotassium phosphate) overnight at 37°C statically.

Subsequently, with the exception of *S. oralis* subsp. *oralis*, we diluted these cultures tenfold in BHI and then used wooden applicator sticks to streak the cultures perpendicularly to the original isolate streak. We used the *S. oralis* subsp. *oralis* culture directly without dilution. We then returned the plates to 37°C and measured the ZOI after overnight incubation. For each set of assays, we inoculated the target strains on a BHI plate without a CF isolate as a control to verify culture viability and purity.

We used ImageJ (117) to measure the ZOI, which we defined as the distance from the original isolate streak to the point at which the target organism began to grow. If a target was not inhibited, we scored the interaction as no inhibition (ZOI = 0 cm). If a target failed to grow on the coculture plate, but grew well on the control plate, we scored the interaction as strong inhibition (ZOI ≥2 cm). We did not score interactions where there was poor growth of the target strain on the control plate or visible contamination of either the isolate or target streak. For consistency, all ZOIs were measured by S.M.M. and spot-checked by R.M.S.

To test whether inhibition differed by target or genus of CF isolate, we performed Chi-square tests of independence and estimated *P*-values by Monte Carlo Simulation (B = 100,000 replicates). To quantify the overall inhibitory activity of individual isolates, we computed the sum inhibition score of each isolate against all eight targets (range: 0 to 24) and then applied Z-score normalization. We limited this analysis to only the 1,364 isolates that were tested successfully against all eight targets. We determined differences in Z-scores across the genera using the Kruskal-Wallis rank-sum test with post-hoc pairwise Wilcoxon rank-sum tests with BH correction for multiple comparisons. We assessed the equality of variance in Z-scores using the Fligner-Killeen test with BH correction for multiple comparisons.

### Characterization of *P. aeruginosa* metabolomes

We inoculated 3 mL BHI with *Pseudomonas* isolates at an initial OD_600_ of 0.0003 and cultured them aerobically at 37°C for 24 h with shaking. Afterwards, we removed cells from 2 mL of culture via centrifugation at 21,130 × g for 20 min at ambient temperature.

We stored 200 *µ*L aliquots at −20°C. We transferred the remaining 1.8 ml of CFS into glass scintillation vials and treated them with three to four drops of chloroform to kill any remaining cells and then evaporated them *in vacuo* using a Biotage V-10 Touch system. We stored the dried CFS at −20°C until processing. To test the CFS for activity, we diluted an overnight culture of *S. aureus* strain PID40 to an OD_600_ of 0.01 and spread 200 *µ*L onto a one-well dish containing 35 mL BHI agar. We then thawed CFS, centrifuged them at 21,130 × g for 5 min at 4°C, spotted 5 *µ*L onto the plate with *S. aureus*, and incubated the culture at 37°C for 24 h before scoring inhibition by identifying ZOIs visually.

We resuspended the dried CFS to 1 mg/mL in MilliQ water, diluted them to 0.1 mg/mL and injected 2 *µ*L for LC-MS/MS analysis. We obtained high-resolution mass spectrometry data using an Agilent 1260 HPLC consisting of a degasser, quaternary pump, autosampler, and column compartment upstream of an Agilent 6545 Q-ToF. Separation was achieved using Kinetex C18 column (50 x 2.1 mm, 100 Å, 2.6 μm, Phenomenex, Torrance, CA) at a flow rate of 0.4 ml/ min held at 30°C. Line C was water with 0.1% (v/v) formic acid and line D was acetonitrile with 0.1% (v/v) formic acid. The column was pre-equilibrated with 97% C/3% D. Upon injection the mobile phase composition was maintained for 0.75 min followed by changing the mobile phase to 10% C/90% D over 9.5 min using a linear gradient. Over the next 0.1 min the mobile phase was changed to 100% D and this composition was held for 3.65 min.

The mobile phase was changed to 97% C/3% D over 0.5 min and held for an additional 1.5 min. The mobile phase held at 97% C/3% D for an additional 3.5 min prior to injection of the next sample. The Agilent Q-ToF mass spectrometer was equipped with an Agilent JetSpray source operated with the following parameters: Auto MS/MS mode, Positive polarity; Gas Temp, 325°C; Drying gas, 10 L/min; Nebulizer, 20 psi; Sheath gas temp, 375°C; Sheath gas flow, 12 L/min; VCap, 4000 V; Nozzle voltage (Expt), 600 V; Fragmentor, 175 V; Skimmer, 65 V; Oct 1 RF Vpp, 750 V; Mass range, 100-3000 *m/z*; Acquisition rate, 10 spectra/s; Time, 100 ms/spectrum. The MS/MS spectra (5 most intense ions) were obtained by data-dependent acquisition with the following parameters by isolating the precursor *m/*z with a medium isolation width (4 *m/*z) summing spectra generated with collision energies of 20, 40, and 60.

We converted raw LC-MS/MS data to .mzXML files using msconvert v3.0.23016 and uploaded them to the XCMS Online server (118). We processed analyses using the Multigroup analysis feature and sample data using the standard HPLC/Q-ToF settings with the following parameters: Feature detection: method, centWave; ppm, 30 minimum peak width, 10; maximum peak width, 60; signal/noise threshold, 6; mzdiff, 0.01; integration method, 2; prefilter peaks, 3; prefiler intensity, 500; noise filter, 0. Retention time correction: method, obiward; profStep, 0.5. Alignment: bw, 5; minifrac, 0.5; mzwld, 0.025; minisamp, 1; max, 100. Statics: statistical test, Kruskal-Wallis non-parametric; performed paired test, view pairs 1; performed post-hoc analysis, true; *p*-value threshold, 0.01; fold-change threshold, 1.5; *P*-value threshold, 0.01; value, into; normalization, none. Annotation: ppm, 5; *m/z* absolute error, 0.015, search for, isotopes. Identification: ppm, 100; adducts, [M+H]^+^; sample biosource, none; pathway ppm deviation, 5; input intensity threshold, 0; significant list *p*-value cutoff, auto. Visualization, EIC width, 200. Miscellaneous: correct mass calibration gaps, no; bypass file sanity check, no.

We performed all downstream analyses on the 468 features that differed significantly across isolates (Kruskal-Wallis *P* ≤ 0.01). For each significant feature, we calculated a multigroup fold change as the ratio of the largest to the smallest non-zero class-mean intensity. We background-corrected the total ion chromatogram (TIC) of every non-blank injection by subtracting the mean TIC of BHI medium blanks, and set any resulting negative values to zero. For hierarchical analysis, we averaged triplicate biological replicates of each isolate to yield a single intensity profile per isolate, log_2_-transformed the profiles, and standardized each feature to a Z-score across isolates. We then clustered profiles by Euclidean distance with complete linkage and estimated support for the clustering by multiscale bootstrap resampling of the features using pvclust (119) with 1,000 bootstrap replicates. To evaluate the contributions of donor and inhibitory phenotype to variation in the metabolome, we corrected the same significant features for medium background by subtracting the mean intensity of the BHI blanks from each feature and setting negative values to zero. We then normalized the blank-corrected intensities to a constant total ion current per sample using total-sum scaling, computed pairwise Bray-Curtis dissimilarities, fit PERMANOVA models, as above, with donor and inhibitory phenotype specified as marginal terms and using 9,999 permutations. We summarized each term using the ω^2^ effect size. We ordinated the samples by principal coordinates analysis (PCoA) and plotted the first two axes.

### Genomic DNA extraction and sequencing of *P. aeruginosa* isolates

We extracted genomic DNA from *P. aeruginosa* isolates adapted a previously reported method (120). Briefly, we washed cell pellets from 3 ml of an overnight culture in BHI broth in 1 ml of Buffer 1 (150 mM NaCl, 10 m M EDTA, 20 mM Tris-HCl, pH 8). We then resuspended the cells in 1 mL of fresh Buffer 1 with 1 mg RNase A and SDS to a final concentration of 1.4% (wt/vol). We incubated the samples at 37°C for 90 min. We then added 0.4 mg of proteinase K and incubated samples at 65°C for 20 min. We then incubated samples at 37°C for 16 h to ensure complete lysis. Subsequently, we extracted and purified genomic DNA used standard phenol-chloroform extraction and precipitation with isopropanol. We submitted genomic DNA to Plasmidsaurus for whole genome sequencing using Oxford Nanopore Technology.

### Genome analyses

We computed ANI among the six isolates and the reference *P. aeruginosa* strain ATCC 10145^T^ (accession: NZ_CP012001.1) using fastANI v1.33 (121). We annotated each genome with Batka v1.8.1 (122) using the following parameters: --genus Pseudomonas --species aeruginosa --complete. We predicted BGCs in each genome with antiSMASH v8.0.4 (123) using the following parameters: --cb-knownclusters --cc-mibig. We identified variants in the isolate genomes using Snippy v4.6.0 (https://github.com/tseemann/snippy). We considered any variant that fell within BGC predicted by antiSMASH to be BGC-associated.

### Statistical analysis and data visualization

We performed all statistical analyses in R with specific packages as described in the preceding sections. We generated graphics using ggplot2 (124) with some cleanup and figure assembly in InkScape.

### Data availability

16S rRNA gene amplicon sequencing data were downloaded from the SRA. Metadata and accession numbers for these runs are provided in **Table S1**. The processed amplicon sequences generated for bacteria isolated for this study are deposited in GenBank under accessions PZ656496 to PZ658092. The genome sequences for *P. aeruginosa* isolates are available under BioProject accession no. PRJNA1491524. The raw mass spectrometry data are deposited at MassIVE under accession: MSV000102457. All other raw data and derived data are available at https://doi.org/10.6084/m9.figshare.32939627. The scripts necessary to replicate this work are available at https://github.com/reedstubbendieck/CF_Interactions.

## Acknowledgements

We thank Elixiva Marcum, Josie Hayes, and Helen Zagloul (Oklahoma State University [OSU]) for assistance with experiments related to this project. We thank Mostafa Elshahed, Sam Miller, and Aymen Yassir (OSU Microbiomics and Culturomics Core Facility) for sequence identification of isolates (P20 GM152333). The Agilent Q-TOF instrument used in this study was purchased with funds from the Division of Research and Innovation and College of Pharmacy at Oregon State University. We thank Paul Lawson (University of Oklahoma) and Matthew Traxler (University of California, Berkeley) for suggestions on the research. We thank Tyrrell Conway (OSU) for comments on the manuscript. We thank all individuals who provided samples for our study, as well as those who contributed samples to the studies we reanalyzed.

This work was supported by the National Institute of General Medical Sciences Centers of Biomedical Research Excellence program (P20 GM152333) and by startup funds from Oklahoma State University to R.M.S., and by internal research funding from Oregon State University to B.P.. S.S. was supported by T32AT010131 (to Drs. Mahmud and van Breemen, Oregon State University). The funders had no role in study design, data collection and interpretation, or the decision to submit the work for publication.

## Author Contributions

**Conceptualization**: Reed M. Stubbendieck

**Data Curation**: Sydney M. Morabbi, Niladri Bhowmik, Shaz Sutherland, Benjamin Philmus, Reed M. Stubbendieck

**Formal analysis**: Sydney M. Morabbi, Niladri Bhowmik, Shaz Sutherland, Benjamin Philmus, Reed M. Stubbendieck

**Funding acquisition**: Benjamin Philmus, Reed M. Stubbendieck

**Investigation**: Sydney M. Morabbi, Niladri Bhowmik, Shaz Sutherland, Evelyn A. Wylie, Reagan S. Decker, Akram Al Daerwish, Mercedes Pérez Pérez, Elizabeth Pascual, Reed M. Stubbendieck

**Resources:** Erika I. Lutter, Reed M. Stubbendieck

**Supervision**: Reed M. Stubbendieck

**Visualization**: Reed M. Stubbendieck

**Writing – original draft:** Reed M. Stubbendieck

## Supplemental Material

**Table S1.** Accessions and metadata for amplicon sequencing samples analyzed in this study.

**Table S2**. Metadata and demographics for donors included in this study.

**Table S3**. Identification of CF Isolates used in this study.

**Table S4.** Genetic variation in *P. aeruginosa* isolates from donor HID07.

**Table S5**. Genetic variation in *P. aeruginosa* isolates from donor HID08.

**Figure S1. CF pathogens bloom to similar relative abundance in the CF airways and to higher levels than oropharyngeal bacteria.** (**A**) Box plots of the genus-level relative abundance of different genera in pathogen-dominated microbiota samples. (**B**) Box plots of the relative of microbiota samples dominated by either CF pathogen-containing or oropharyngeal genera. The upper and lower bounds of the box plots indicate the 75th and 25th percentiles, respectively. The horizontal black bars indicate the medians, and the notches represent the 95% confidence intervals of the medians. The whiskers extend from the bounds of the box to the largest and smallest values that are no further than ±1.5× the interquartile range (IQR). In **A**, Genera that share letters are not significantly different (Benjamini-Hochberg-adjusted *P* > 0.05).

**Figure S2. Composition of the CF airway microbiota does not vary based on age, lung function, or sex.** Each point in the nonmetric multidimensional scaling (NMDS) plots represents the genus-level bacterial community sequenced from one individual. The points are colored based on age (**A**), lung function category (**B**), or sex (**C**), as indicated by the respective keys below each plot. The ellipses represent the 95% confidence regions for each group. The stress value for each NMDS is shown in the lower right of the panel.

**Figure S3. *Pseudomonas* is the most common pathogen that dominates the airways of pwCF.** Plot showing the relationship between the Berger-Parker (BP) Dominance and lung function. Each microbiota sample is indicated with a point. Points were jittered to avoid overplotting. The color of each point indicates the most abundant genus, as indicated by the key below. Points to the right of the dashed line are dominated by a single genus. The red line indicates the line of best fit, and each of the 100 light right lines represents a bootstrap replicate of the fit.

**Figure S4. Variable inhibitory activity of *Pseudomonas* isolates from the same donor and timepoint.** Plot showing the Z-score normalized inhibition of *Pseudomonas* isolates from different donors tested against all respiratory target strains that possessed variable activity (Z-score spread >2). Each point represents one isolate. Each point represents one isolate. Donor IDs in bold indicate hosts from which isolates were selected for genome sequencing and metabolomics, indicated in gold. The upper and lower bounds of the box plots indicate the 75th and 25th percentiles, respectively. The horizontal black bars indicate the medians. The whiskers extend from the bounds of the box to the largest and smallest values that are no further than ±1.5× the IQR.

**Figure S5. Donor source structures *Pseudomonas* metabolomes.** (**A**) Inhibitory activity of cell-free supernatants (CFS) from *Pseudomonas* isolates against *S. aureus*. All CFS are grouped based on the donor. Images are representative of biological triplicates. (**B**) Principal coordinate analysis plot based on Bray-Curtis dissimilarity compared from 468 molecular features that varied across six *P. aeruginosa* isolates in biological triplicate after subtraction of the averaged media blank. Each point represents the metabolome profile of a single *P. aeruginosa* isolate. The color and shape of the points represent the donor identity (HID) and inhibitory activity of the isolate, respectively. The solid and dashed ellipses represent the *t*-distribution spread for HID and inhibitory activity, respectively.

## Notes

### Competing Interest Statement

The authors have declared no competing interest.

### Summary of Updates

Added supplementary material that was not uploaded by publisher. Fixed typos.

