## Supplementary figures and images for "Differential Metabolite Production Underlies Disruption of the Cystic Fibrosis Airway Microbiota by Pathogens"

### Fig. S1

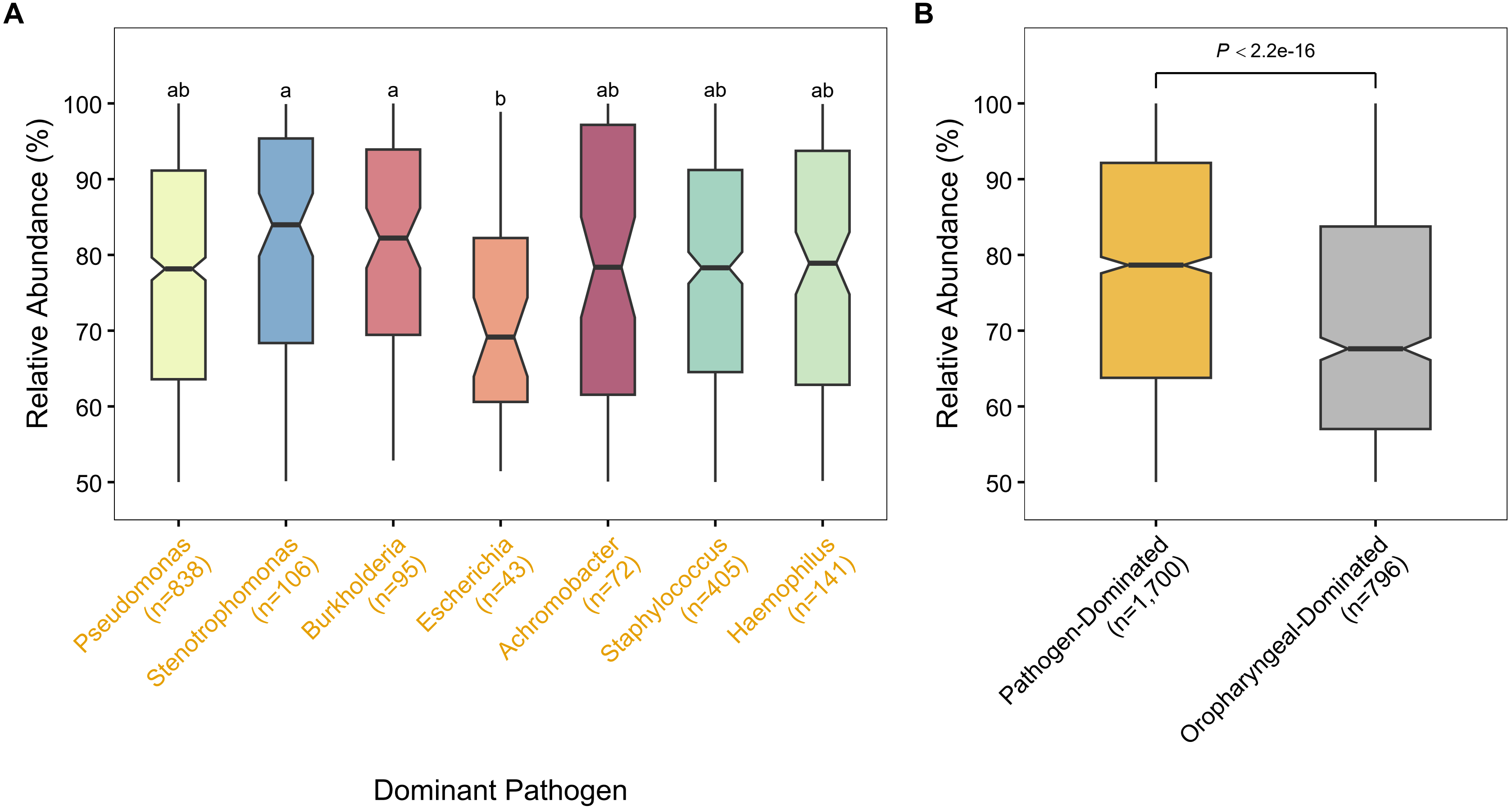

### Fig. S2

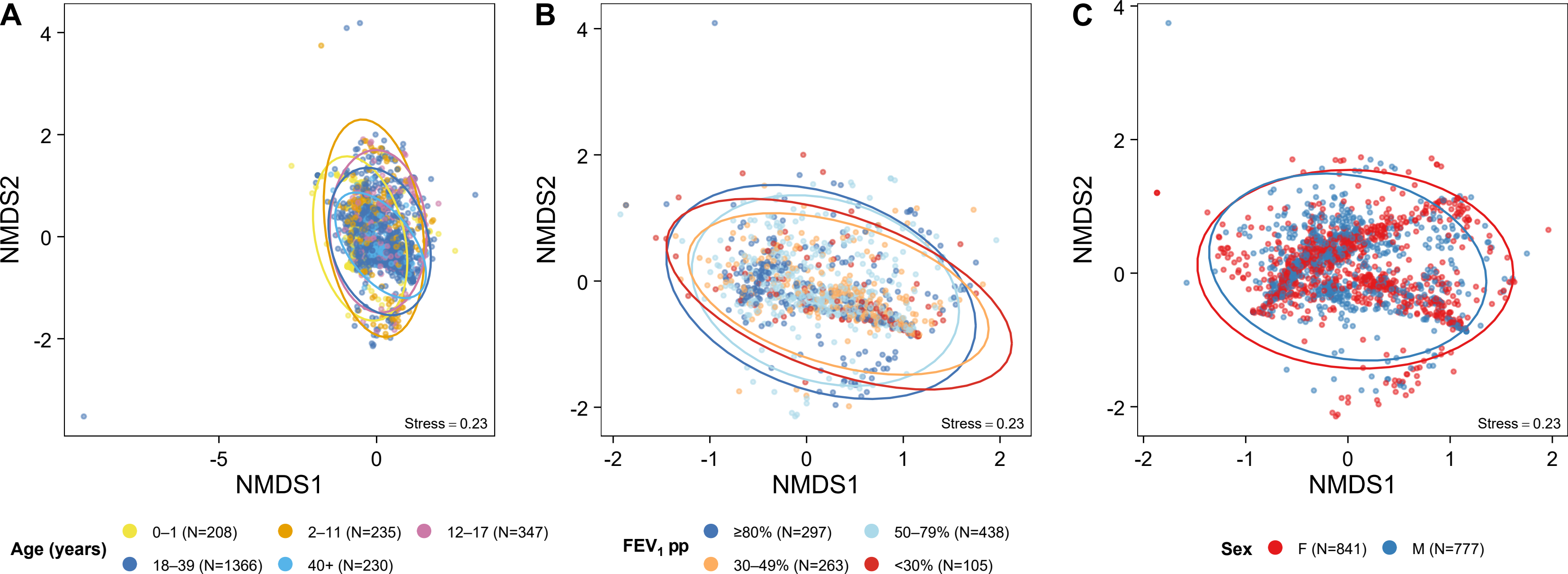

### Fig. S3

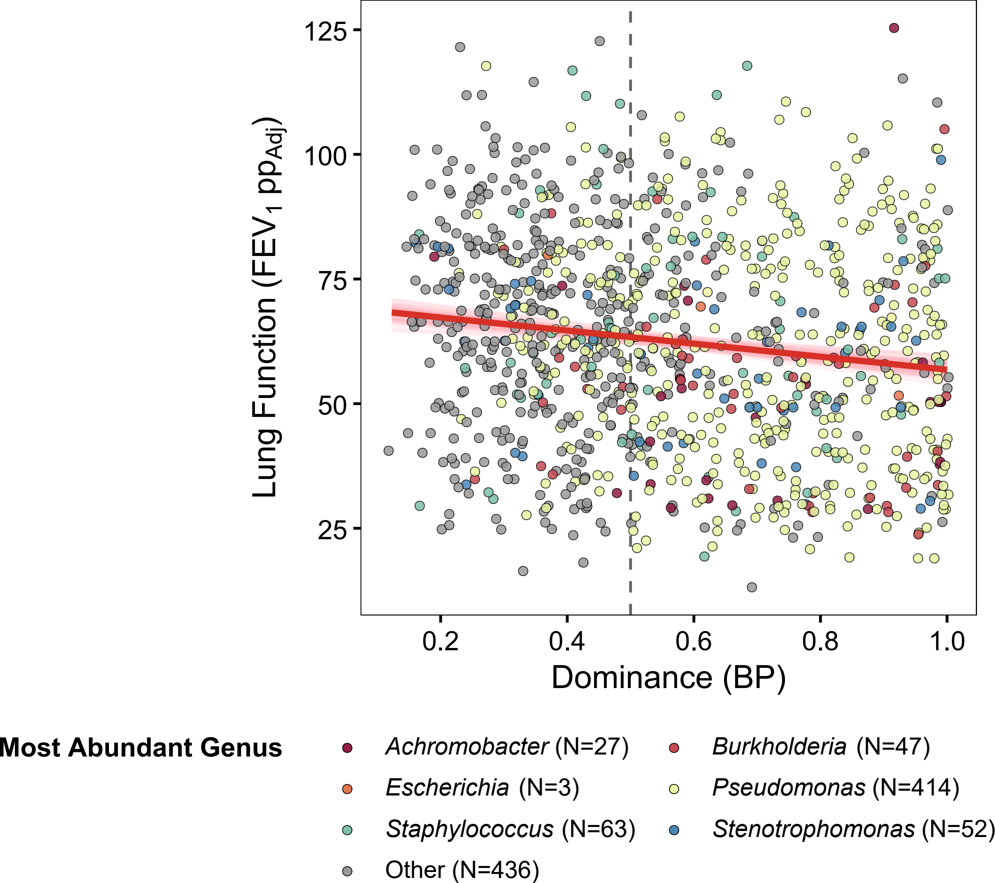

### Fig. S4

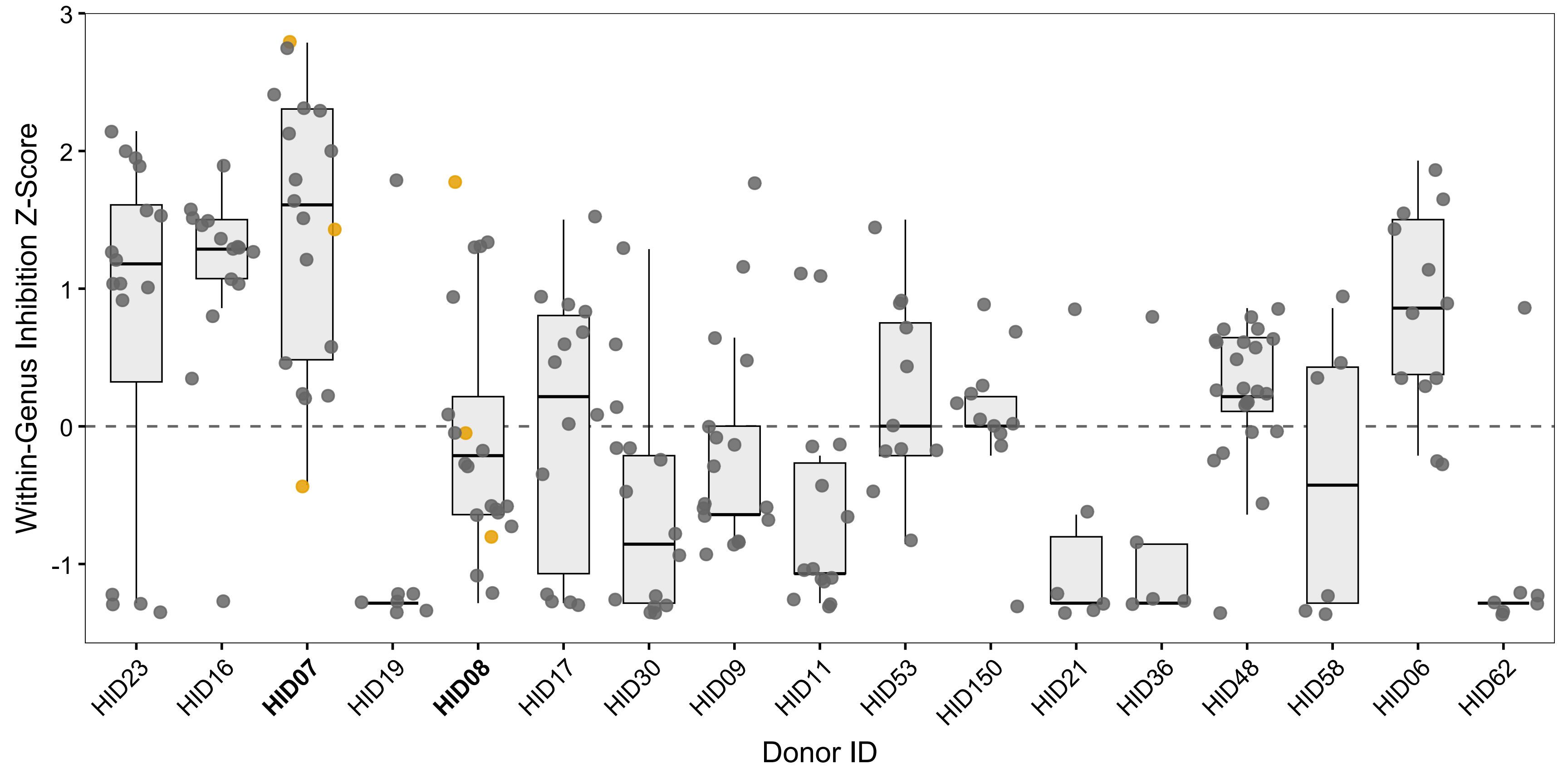

### Fig. S5

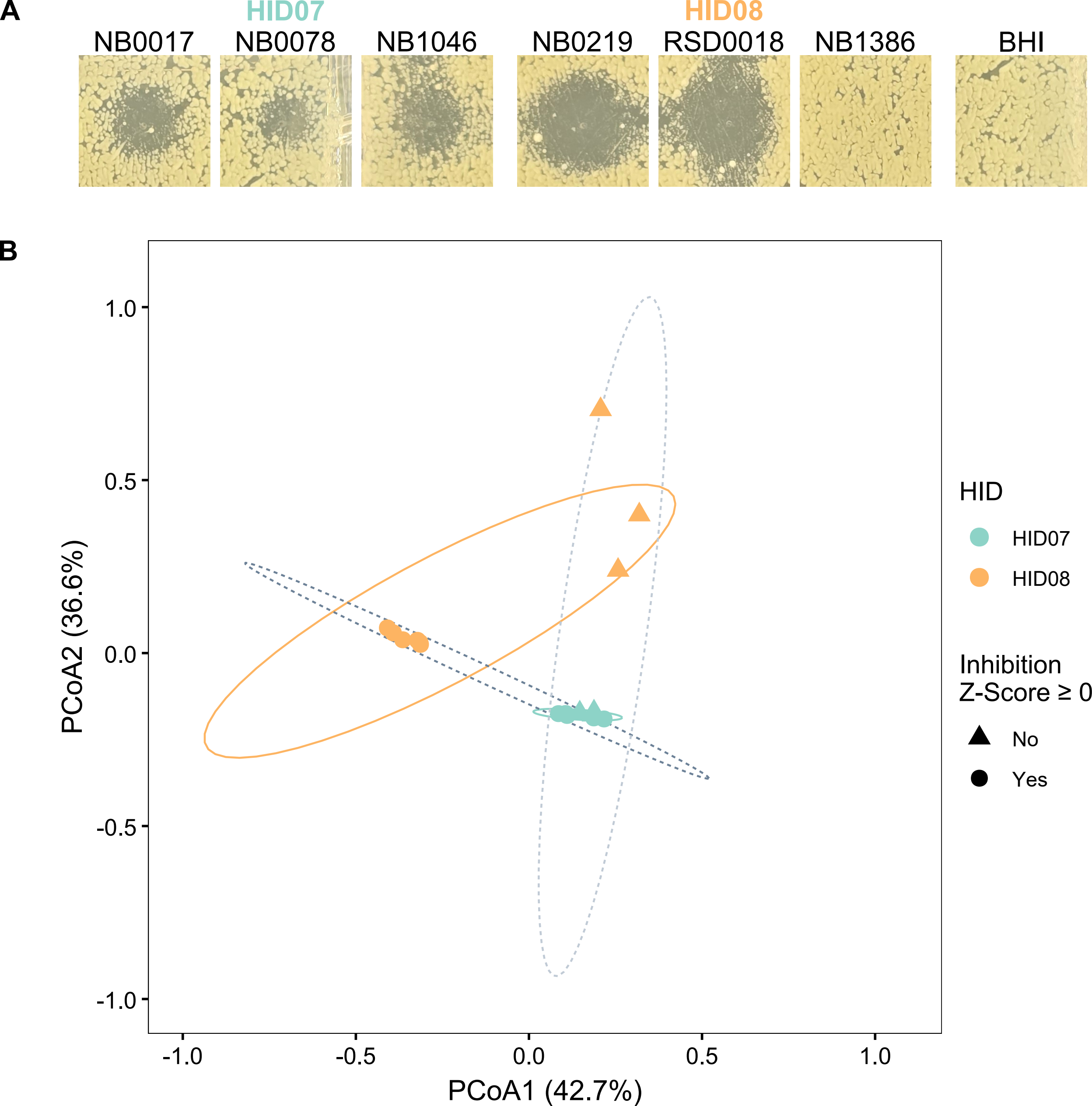
